# Loss of FSTL1-expressing adipocyte progenitors drives the age-related involution of brown adipose tissue

**DOI:** 10.1101/2020.05.14.096990

**Authors:** Zan Huang, Zengdi Zhang, Ryan Heck, Ping Hu, Hezkiel Nanda, Kaiqun Ren, Zequn Sun, Alessandro Bartolomucci, Yan Gao, Dongjun Chung, Weiyun Zhu, Steven Shen, Hai-Bin Ruan

**Author notes:** These authors contributed equally (Zan Huang, Zengdi Zhang). Correspondence: H.-B. R. or Z. H. Lead contact: H.-B. R.

## Abstract

In humans, brown adipose tissue (BAT) undergoes progressive involution or atrophy with increasing age, as manifested by decreased prevalence and mass, transformation to white adipose tissue (WAT), and reduction in thermogenic activity. This involution process cannot be fully recapitulated in rodent models and thus underlying cellular mechanisms are poorly understood. Here, we show that the interscapular BAT (iBAT) in rabbits involutes rapidly in early life, similarly to that in humans. The transcriptomic remodeling and identity switch of mature adipocytes are accompanied with the loss of brown adipogenic competence of their precursor cells. Through single-cell RNA sequencing, we surveyed the heterogenous populations of mesenchymal cells within the stromal vascular fraction of rabbit and human iBAT. An analogous *FSTL1*_high_ population of brown adipocyte progenitors exists in both species while gradually disappear during iBAT involution in rabbits. In mice, FSTL1 is highly expressed by adipocyte progenitors in iBAT and genetic deletion of FSTL1 causes defective WNT signaling and iBAT atrophy in neonates. Our results underscore the BAT-intrinsic contribution from *FSTL1*^high^ progenitors to age-related tissue involution and point to a potential therapeutic approach for obesity and its comorbidities.

**HIGHLIGHTS:**

1. Rabbit BAT irreversibly transforms to WAT before puberty.
2. iBAT adipocyte progenitors reprogram transcriptome and lose brown adipogenic ability.
3. Comparable *FSTL1*^high^ brown adipocyte progenitors exist in rabbit and human iBAT.
4. Loss of FSTL1 in brown adipocyte progenitors causes iBAT atrophy in mice.

## INTRODUCTION

Obesity is a strong predisposing factor for many diseases, including hypertension, high cholesterol, diabetes, and cardiovascular diseases. The thermogenic brown adipose tissue (BAT) not only dissipates chemical energy as heat to maintain body temperature and energy balance (Cannon and Nedergaard, 2004; Rosen and Spiegelman, 2014; Yang and Ruan, 2015), but also serves as a metabolic sink for glucose, fatty acids, and branched-chain amino acids to improve metabolic health (Kajimura et al., 2015; Wallace et al., 2018; Maurer et al., 2019; Yoneshiro et al., 2019). Therefore, BAT has been increasingly acknowledged as a potential therapeutic target for obesity and related disorders.

In humans, there seem to be two spatiotemporally distinct phases of BAT ontogeny. BAT first appears mainly in interscapular and perirenal regions during mid-gestation (Lidell, 2019). While maximally recruited in neonates, the interscapular BAT (iBAT) rapidly atrophies till undetectable in adults (Heaton, 1972; Sidossis and Kajimura, 2015). Perirenal BAT also steadily transforms to white adipose tissue (WAT) with increasing age (Tanuma et al., 1975). The second developmental phase occurs postnatally and gives rise to anatomically dispersed BAT in cervical, supraclavicular, and axillary areas in adult humans (Rogers, 2015). The existence of metabolically active BAT in adult humans was rediscovered by [^18^F]-Fluorodeoxyglucose-PET/CT imaging and subsequent histological and molecular characterization (Cypess et al., 2009; Saito et al., 2009; van Marken Lichtenbelt et al., 2009; Virtanen et al., 2009; Zingaretti et al., 2009). Despite that the absolute amount and energy-burning activity of human BAT have not yet been ascertained (Lee et al., 2013), PET/CT studies have revealed a consensus that the prevalence of cervical-supraclavicular BAT peaks in adolescence and then reduces to less than 5% in middle-aged men and women (Rogers, 2015). Cold recruits BAT, even in subjects having no existing metabolically active BAT (Saito et al., 2009; van Marken Lichtenbelt and Schrauwen, 2011; Yoneshiro et al., 2011). But such recruitment reportedly happens only in young adults and there is no evidence that BAT can be efficiently recruited in aged people with no detectable BAT. The age-related atrophy of BAT may contribute to the decline of metabolic health in elderly; however, cellular and molecular mechanisms governing the involution process remain enigmatic.

Mice possess human-like BAT and beige adipose depots in anatomically comparable regions (Mo et al., 2017; Zhang et al., 2018). Using mouse models, extensive knowledge has been gained on the lineage development, transcriptional regulation, and systemic function of thermogenic adipocytes (Harms and Seale, 2013; Sanchez-Gurmaches et al., 2016; Scheele and Wolfrum, 2020). It is well accepted that the dorsal-anterior-located interscapular fat depots predominately arise from the *Pax3^+^/Meox1^+^/Myf5^+^* mesenchymal lineage (Sebo and Rodeheffer, 2019). Within the adipose tissue, adipocyte stem/progenitor cells and committed preadipocytes reside in the stromal vascular fraction (SVF) and normally express lineage-negative markers including PDGFRα, DLK1, CD34, SCA1, CD29, and CD24 (Rodeheffer et al., 2008; Schulz et al., 2011; Berry and Rodeheffer, 2013). Profiling of the SVF cells from WAT depots at the single-cell level has revealed a developmental hierarchy of white adipocyte precursors (Burl et al., 2018; Hepler et al., 2018; Schwalie et al., 2018; Cho et al., 2019; Merrick et al., 2019; Rajbhandari et al., 2019). However, the cellular heterogeneity and differentiation trajectory of brown adipose precursors have not been defined yet.

Compared to that in humans, there is limited involution or atrophy of BAT associated with life history in mice. iBAT in rodents indeed undergoes age-dependent remodeling that is characterized by tissue hypertrophy and whitening, reduced expression of the mitochondrial uncoupling protein 1 (UCP1), and thermogenic dysfunction (Horan et al., 1988; McDonald et al., 1988; Florez-Duquet et al., 1998). However, iBAT adipocytes remain as the progeny of *Myf5^+^* cells (Sanchez-Gurmaches and Guertin, 2014), and can be immediately recruited by external cues such as cold and sympathetic activation in old mice, although to a slightly less extent (Sellayah and Sikder, 2014; Goncalves et al., 2017; Tajima et al., 2019a). At thermoneutral conditions, iBAT in mice also retains its browning capacity, despite of reduced expression of thermogenic genes (Cui et al., 2016; Roh et al., 2018). These observations not only indicate that mice are inadequate in modeling the involution process of iBAT but also suggest that cell-intrinsic mechanisms exist to control the divergent aging program of iBAT across species. Here, we establish rabbits as an alternative model to investigate BAT involution in humans. Rabbit iBAT irreversibly transforms to WAT before puberty, at the similar developmental stage when human iBAT disappears. Adipocyte precursors in iBAT of rabbits reprogram the transcriptome and lose their capability to be differentiated into mature brown adipocytes. Single-cell RNA sequencing (scRNA-seq) of SVF cells from iBAT identifies comparable *FSTL1*^high^ cells as brown adipocyte progenitors in both neonatal rabbits and fetal humans. The constitutive presence of iBAT in mice is associated with the high-level expression of FSTL1 in adipocyte progenitors. Genetic deletion of FSTL1 in brown adipocyte progenitors causes iBAT atrophy in mice. In future, preserving or regenerating *FSTL1*^high^ brown adipocyte progenitors may prevent BAT involution or rejuvenate aged BAT in elderly, thereby promoting metabolic fitness.

## RESULTS

### Progressive BAT whitening in young rabbits

To characterize BAT involution in rabbits, we collected interscapular, subscapular, suprascapular, cervical, supraclavicular, axillary, and perirenal fat depots from New Zealand White rabbits at the ages of 1 day, 3, 6, and 12 weeks. Histology showed that most depots had the typical morphology of BAT in neonatal rabbits, except that axillary fat was a mixture of multi-and uni-locular adipocytes and most cells in the perirenal depot were unilocular fat cells (**Figure 1A**). Tissue whitening, adipocyte hypertrophy, and loss of UCP1 expression could be readily seen starting at 3 weeks of age (**Figure 1A, B**). Interscapular, subscapular, suprascapular, and cervical depots were all converted to WAT-like tissues at 12 weeks of age. Supraclavicular and axillary BAT whitened at a slightly quicker pace, containing mostly unilocular white adipocytes at 6 weeks of age. Perirenal fat was whitened as early as 3 weeks. These data show that, though at different rates, all rabbit BAT depots that we have examined undergo rapid whitening before puberty, similar to interscapular BAT (iBAT) involution observed in humans.

**Figure 1.**
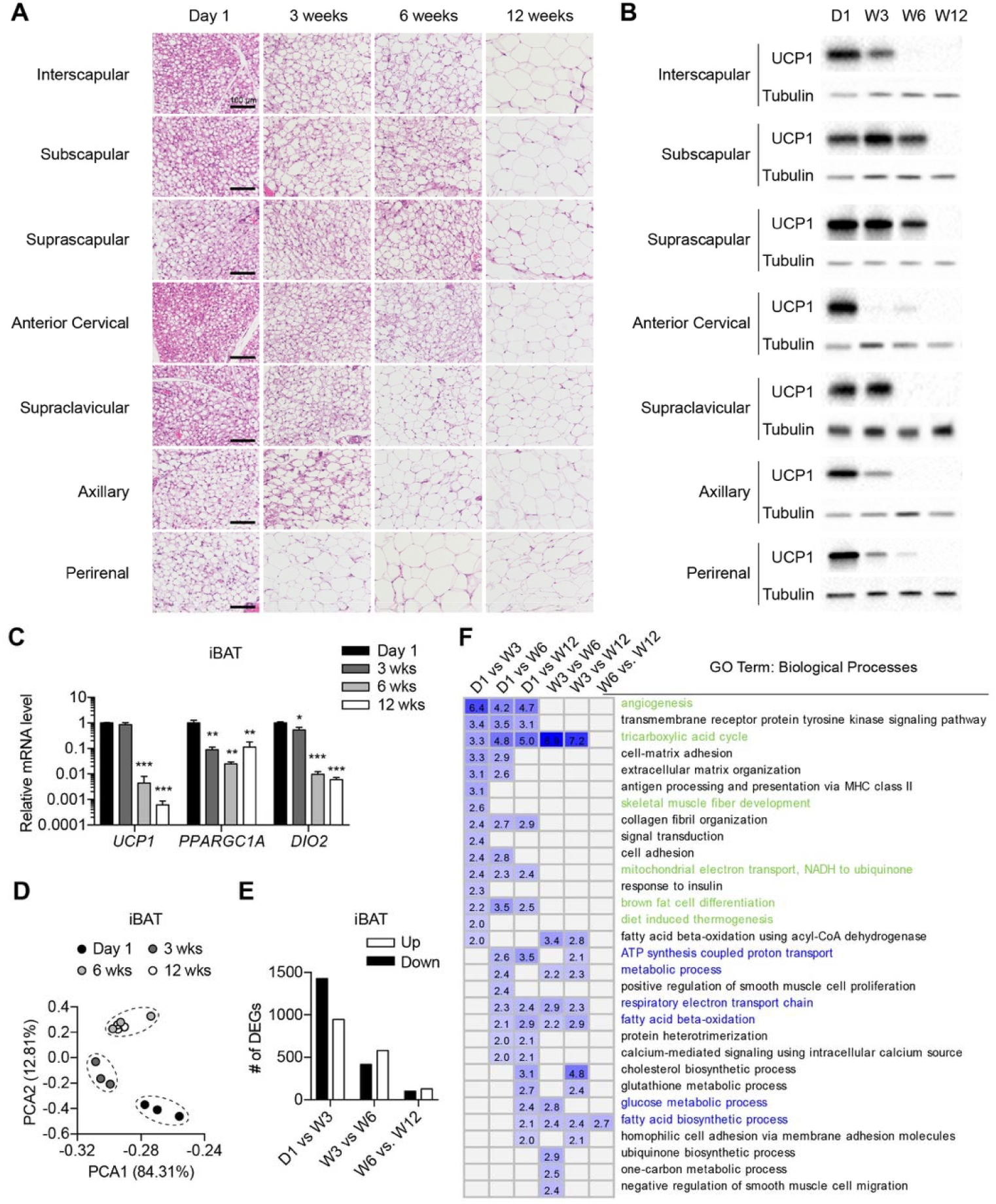
Histological and transcriptional remodeling of rabbit BAT. (A, B) Representative H&E images (A) and UCP1 protein expression (B) of BAT depots from rabbits at different ages (n = 3). Scale bar = 100 μm. (C) Expression of *Ucp1, Ppargac1,* and *Dio2* transcripts in total iBAT (n = 3). (D) Principal component analysis (PCA) plot of rabbit iBAT at different ages after RNA-seq. (E) Numbers of differently expressed genes between iBAT samples form indicated ages. (F) Gene Ontology (GO) classification and functional enrichment by DAVID. P values (-log10) were shown as heatmap. Data are presented as mean ± SEM. *, P < 0.05; **, P < 0.01; ***, P < 0.001 by one-way ANOVA followed with Tukey’s multiple comparison (C).

To verify morphological observations, we focused on iBAT and examined thermogenic gene expression. *UCP1* mRNA levels in total rabbit iBAT were comparable between day 1 and week 3, but markedly reduced more than 500 folds at 6 weeks of age and 5000 folds at 12 weeks of age (**Figure 1C**). The extent of reduction of *UCP1* mRNA in rabbits was much greater than that in mice transitioned from room temperature to thermoneutrality (Cui et al., 2016; Roh et al., 2018). Upstream regulators of *UCP1* including *PPARGC1A* and *DIO2* started to reduce their expression as early as 3 weeks (**Figure 1C**). We then performed RNA sequencing of all transcripts from total iBAT. Principal component and gene expression correlation analyses showed a clear separation of iBAT from neonatal and 3-week-old rabbits, while 6- and 12-week samples were clustered together (**Figure 1D** and **Figure S1A, B**). The majority of changes in gene expression happened at 3 weeks of age and there were much fewer differentially expressed genes at 6 and 12 weeks of age (**Figure 1E**). These data suggest that the transcriptomic reprogramming in iBAT occurs early in life, even before the reduction of *UCP1* transcripts. Hierarchical clustering and pathway analysis showed that the early differentially-expressed genes were involved in processes including angiogenesis, tricarboxylic acid (TCA) cycle, skeletal muscle development, mitochondrial electron transport, and brown fat differentiation and thermogenesis (**Figure 1F** and **Figure S1C**), which were all associated with BAT function. On the other hand, late-changing genes controlled metabolic processes like ATP synthesis, fatty acid oxidation and biosynthesis, and glucose metabolism (**Figure 1F**), probably representing the metabolic adaptation to tissue whitening. Collectively, rabbits are a suitable model to investigate age-related involution of BAT.

### Switch of adipocyte identity during BAT involution

Were those unilocular and UCP1-negative adipocytes found in regressed rabbit iBAT white adipocytes or “dormant”, inactive brown adipocytes? To determine the cellular identity of fat cells, we first examined the expression of different adipocyte marker genes that have been identified in mice and humans. Brown adipocyte markers were all dramatically downregulated from Day 1 to Week 12 in rabbit iBAT (**Figure S1D**). The levels of beige markers including *TBX1* and *CD137* were relatively stable (Roh et al., 2018), while expression of white markers gradually increased over time (**Figure S1D**). Sympathetic nerve activity controls lipolysis and thermogenesis in BAT, and we found a substantial reduction in the expression of β1-3 *(ADRB1-3)* and α1A, B *(ADRA1A* and *ADRA1B)* adrenergic receptors and an upregulation of α1D *(ADRA1D)* and α2 *(ADRA2A* and *ADRA2B)* receptors in the rabbit iBAT (**Figure S1E**), suggesting an active, intrinsic change in the adrenergic signaling.

We then sought to determine if whitened iBAT can be remodeled by cold. Adult rabbits were purchased from a non-climate-controlled rabbitry in Wisconsin during summer (August 2017 with an average temperature of 21°C) and early Spring (March 2018 with an average temperature of 0 °C). Cold environment did not prevent the whitening and loss of UCP1 in adult iBAT (**Figure 2A, B**). It has been shown in mice that cold stimulates beige adipogenesis from perivascular progenitors while β3-adrenergic receptor activation induces the conversion from mature white to beige adipocytes (Jiang et al., 2017; Chen et al., 2019). We treated adult rabbits for 2 weeks with Mirabegron, a β3-adrenergic receptor agonist that also works in rabbits (Calmasini et al., 2015). Mirabegron did recruit a few multilocular “brown-like” adipocytes in iBAT and induce *UCP1* gene expression with high variability (**Figure 2C, D**). These data suggest that most fat cells in adult rabbits were not “dormant” brown adipocytes, but white or beige adipocytes. Future development of genetic rabbit models and fate-mapping experiments are desired to determine whether brown adipocytes are transdifferentiated into white/beige adipocytes or they are replaced by new white fat cells differentiated from their precursors.

**Figure 2.**
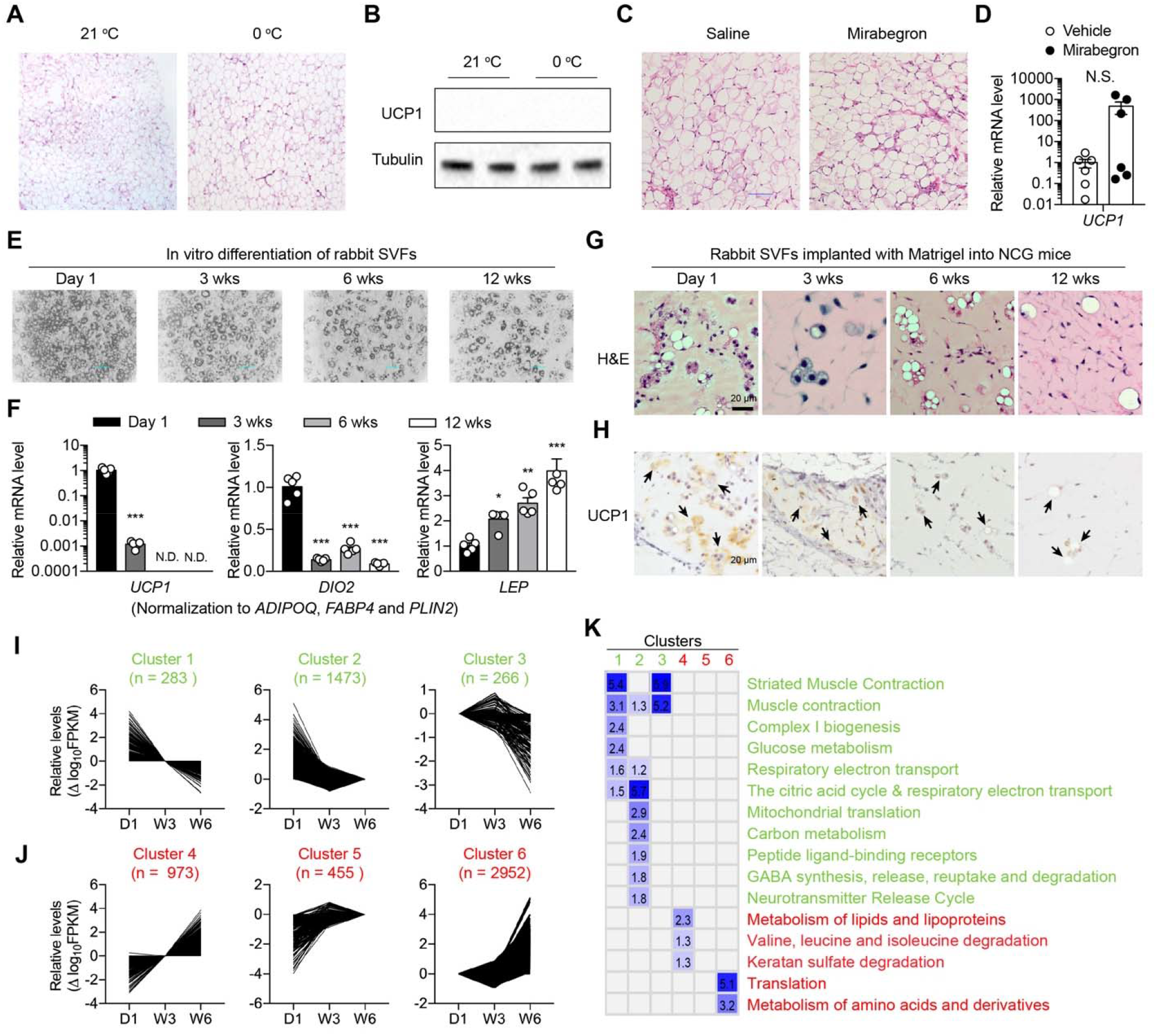
Identify switch of BAT adipocytes and SVF progenitors. (A, B) Representative H&E images (A) and UCP1 protein expression (B) of iBAT depots from adult rabbits collected at summer or early spring (n = 2). (C, D) Adult rabbits were treated with saline or Mirabegron for 2 weeks (n = 6). iBAT histology and *UCP1* gene expression were determined. N.S., not significant. (E, F) iBAT SVF cells from rabbits at different ages were differentiated into adipocytes. (E) Lipid deposition in differentiated cells. (F) Relative gene expression normalized to geometric mean of *ADIPOQ, FABP4,* and *PLIN2* (n = 5). (G, H) SVF cells from periscapular BAT of rabbits at different ages were isolated, mixed with Matrigel, and implanted into NCG mice (n = 3). Representative H&E (G) and UCP1 immunohistochemistry (H) of implants are shown. Arrows indicate adipocytes. (I, J) Clusters of genes showing down-regulation (I) and up-regulation (J) from day 1 to week 6. (K) GO biological processes enriched in each cluster. P values (-log10) are shown as heatmap. Data are presented as mean ± SEM. *, P < 0.05; **, P < 0.01; ***, P < 0.001 by one-way ANOVA followed with Tukey’s multiple comparison.

### Age-related loss of the brown adipogenic ability of adipocyte progenitors

One potential contributing factor for BAT involution could be the impairment in the browning capacity of adipocyte progenitor cells. In vitro, the stromal vascular fraction (SVF) cells from iBAT of neonatal rabbits that contain adipocyte precursors could be differentiated into *UCP1*-expressing brown adipocytes (Cambon et al., 1998) (**Figure 2E, F**). As rabbits grow, while no changes in cell proliferation were observed (**Figure S2A**), SVF cells from 3 to 12-week-old rabbits displayed slightly reduced adipogenic differentiation as shown by less lipid deposition (**Figure 2E**) and lower expression levels of the *ADIPOQ, FABP4,* and *PLIN2* genes (**Figure S2B**). Nevertheless, the expression of thermogenic *UCP1* (about 1000-fold lower at 3-week and not detectable at 6-and 12-week) and *DIO2* was more drastically downregulated, either normalized to the housekeeping gene *RPLP0* (**Figure S2C, D**) or normalized to the geometric mean of *ADIPOQ, FABP4,* and *PLIN2* genes (**Figure 2E**). Conversely, the expression of white adipocyte-specific *LEP* gene in differentiated adipocytes was positively correlated with rabbit age (**Figure 2E**).

To further determine the brown adipogenic potential of adipocyte progenitors in vivo, we isolated SVF cells from inter- and sub-scapular BAT (iBAT and sBAT) of rabbits at different ages, mixed and resuspended in Matrigel, and implanted into the dorsal region of immune-deficient *NOD-Prkdc^em26Cd52^ Il2rg^em26Cd22^/NiuCrl* (NCG) mice (Min et al., 2016). H&E staining and UCP1 immunohistochemistry showed that SVF cells from neonatal rabbits could form largely multilocular and UCP1-positive brown adipocytes (**Figure 2G, H**). However, in the Matrigel implants derived from older SVF cells, unilocular and UCP1-low and -negative adipocytes gradually emerged and finally dominated (**Figure 2G, H**). From these data, we conclude that there is an age-related reduction in brown adipogenic ability of periscapular SVF cells in rabbits.

Correlated with the reduced ability to be differentiated into brown adipocytes, there was an age-associated transcriptional reprograming in SVF cells from rabbit iBAT and sBAT (**Figure S2E, F)**. RNA-seq of SVF cells revealed a large number of differentially expressed genes from day 1 to 3 weeks of age (**Figure S2G, H**). Much fewer changes in gene expression were observed between week 3 and 6 and another wave of transcriptional change was observed at week 12 (**Figure S2C, D**). Though iBAT and sBAT depots were not fully whitened until rabbits were 12 weeks old (**Figure 1A**), the transcriptional remodeling of total iBAT was observed mainly at week 3 and 6 (**Figure 1D, E**). We thus suspected that the molecular changes in brown adipocyte progenitors that potentially contribute to BAT involution should occur before week 6, and the differentially expressed genes in SVF cells at week 12 could be adaptive responses to tissue whitening (**Figure 1F**). Therefore, transcriptional data of SVF cells excluding the 12-week age point were subjected for longitudinal analyses. Differentially expressed genes in iBAT and sBAT SVF cells were combined for the clustering analysis. Co-expressed genes could be mainly grouped into six clusters. Clusters 1 to 3 showed down-regulation with distinct patterns: continued down-regulation, early down-regulation, and late down-regulation (**Figure 2I**). And clusters 4 to 6 represented genes had continued up-regulation, early up-regulation, and late upregulation, respectively (**Figure 2J**). Pathway analysis showed that down-regulated genes were enriched for muscle contraction and mitochondrial function, while up-regulated genes were involved in translation and metabolism of lipid and amino acid (**Figure 2K**). Collectively, these data demonstrate that periscapular BAT involution in rabbits is associated with identify switch of mature adipocytes and transcriptional reprograming of their progenitors.

### Dynamic transcriptional landscape of rabbit BAT stromal cells at the single-cell level

The SVF of BAT contains a diversity of cell populations such as mesenchymal progenitors, preadipocytes, fibroblasts, endothelial cells, and immune cells. To understand the developmental heterogeneity and dynamics of the BAT stromal cells, we then performed singlecell RNA sequencing (scRNA-seq) of freshly isolated iBAT SVF cells from rabbits at the age of 1 day, 3 weeks, and 12 weeks using the 10X Genomics platform. After excluding *CD45^+^* hematopoietic cells, total of 8719 cells (1488, 5508, and 1723 cells from day 1, 3-week, and 12-week rabbits, respectively) were analyzed (**Table S1**). Unsupervised clustering of the gene expression profiles identified 7 cell populations (**Figure 3A** and **Table S2**). Canonical adipose mesenchymal progenitor markers including *WNT2, DLK1,* and *THY1 (CD90)* were expressed predominantly in group 1, 0, and 6 (**Figure 3B** and **Figure S3A**). Adipocyte identity genes, including *FABP4, LPL, PPARG,* and *APOD,* were specifically enriched in group 2 and 4 cells (**Figure 3C** and **Figure S3B**). By comparing differentially expressed genes between groups 1/0/6 and 2/4 and then identifying potential upstream regulators (**Table S3**), we found that groups 1/0/6 highly expressed genes that are activated by anti-adipogenic TGFB1, thus representing adipocyte progenitors (**Figure S3C**). While actively expressed genes in groups 2/4 were controlled by PPARG, PPARA, and CEBPB (**Figure S3C**), indicating that these cells were “committed preadipocytes”. Previously identified mesenchymal markers in mice and humans, such as *PDGFRA, PDGFRB,* and *CD34* were expressed by all cell populations, with *PDGFRA* and *PDGFRB* have a trend of increased expression in preadipocytes (**Figure S3A**). Furthermore, by searching for marker genes that encode cell surface proteins, we identified *ADGRD1 (GPR133)* and *SFRP4* as novel markers for adipocyte progenitors (**Figure 3A**) and *VCAM1 (CD106)* and *NRP1 (CD304)* as preadipocyte markers (**Figure 3B**), some of which were used later for flow cytometric analyses of these populations in mice.

**Figure 3.**
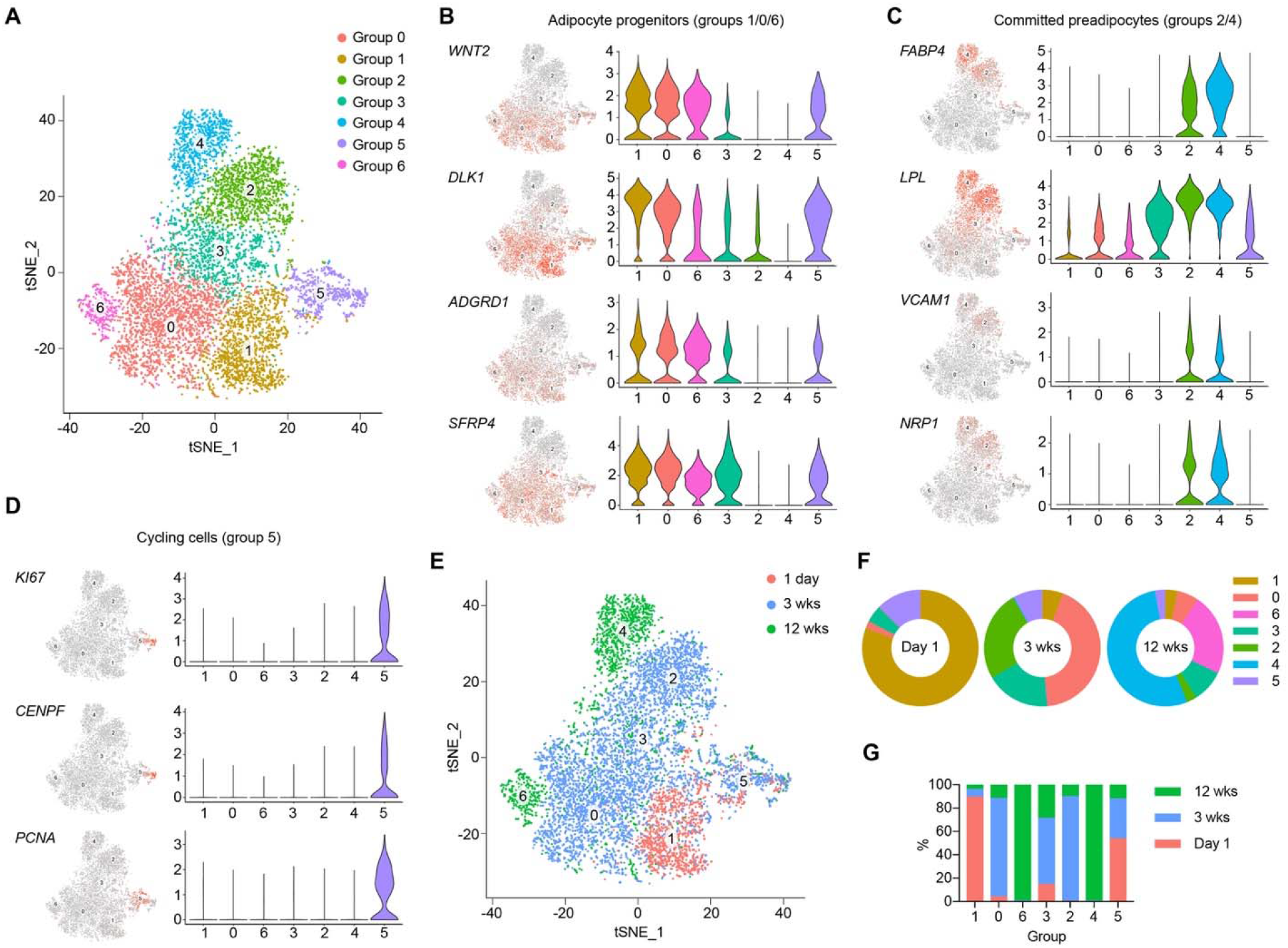
Transcriptional dynamics of rabbit iBAT SVF cells at the single-cell level. (A) tSNE plot showing 7 distinct groups of cells clustered from pooled *PTPRC* (*CD45*)-negative iBAT SVFs of 1-day, 3-week, and 12-week old rabbits. (B-D) tSNE and violin plots showing the expression levels and distribution of marker genes for adipocyte progenitors (B), committed preadipocytes (C), and cycling cells (D). (E) tSNE plot of iBAT SVF cells segregated by age. (F) Cellular composition of SVF cells at different ages. (G) Distribution of different-aged SVF cells within each group.

Group 3 cells started to reduce the expression of mesenchymal markers while began to express preadipocyte genes (**Figure 3A-C** and **Figure S3A, B**). Moreover, group 3-enriched genes such as *FN1* and *OLFML2B* have been shown to be transiently expressed during adipogenesis (Ojima et al., 2016), indicating that group 3 were intermediate cells differentiating from progenitors to committed preadipocytes (**Figure S3D**). Group 5 cells were actively cycling cells as genes involved in cell cycle control such as *MKI67, CENPF,* and *PCNA* were exclusively expressed within this group (**Figure 3D**).

SVF cells from rabbits at different ages were readily separated, pointing to age-related reprograming of the transcriptome at the single-cell level (**Figure 3E**). The majority of day 1 SVF cells were group 1 adipocyte progenitors (80.5%) and cycling cells (12.6%), with very few cells were differentiating or committed preadipocytes (**Figure 3F** and **Table S1**). 3-week SVF cells were largely composed of group 0 adipocyte progenitors (42.7%), group 3 differentiating precursors (17.8%), and group 2 committed preadipocytes (25.2%). At 12 weeks of age, group 6 adipocyte progenitors (22.6%) and group 4 committed preadipocytes (53.2%) were dominant. After normalizing to cell number, we could observe a gradual transition of adipocyte progenitors from group 1 to group 0 and then to group 6 as rabbits grew older (**Figure 3G** and **Table S1**). Almost no committed preadipocytes (i.e. groups 2 and 4) were found in neonatal rabbits. In addition, as rabbits age, there were less and less group 5 cycling cells (**Figure 3G** and **Table S1**), indicating precursor cells transitioning from self-renewal to differentiation. Collectively, these data demonstrate that iBAT involution in rabbits is associated with cellular compositionshifting and transcriptional reprograming of the adipose progenitors/preadipocytes.

### Identification of conserved markers for brown adipose progenitors between rabbits and humans

Adipocyte progenitors in older rabbits gradually lost their brown adipogenic ability (**Figure 2G, H**) (Cambon et al., 1998) and shifted their transcriptomic landscape (**Figure 2I-K and Figure 3**). We thus reasoned that Group 1 cells revealed by scRNA-seq were rabbit brown adipocyte progenitors. We listed both positively and negatively enriched genes of group 1 cells and performed pathway analysis. Top activated pathways included EIF2 and mTOR pathways, while LXR/RXR and PPAR signaling were inhibited (**Figure S4A**). This was consistent with previous findings that mTOR and protein translation are essential for BAT development and thermogenesis (Ye et al., 2019).

iBAT in both rabbits and humans undergoes rapid involution after birth. An important question is whether analogous populations of adipocyte progenitors exist in rabbits and humans. To address this, we isolated SVF cells from iBAT of a human fetus at the gestational age of 24 weeks after an induced abortion and subjected them to scRNA-seq analysis. We identified four groups (0, 1, 3, and 2) of *PDGFRa* adipogenic lineage cells, as well as pericytes, endothelial cells, lymphatic endothelial cells, hematopoietic/immune cells, and actively cycling cells (**Figure 4A, Figure S4B, C,** and **Table S4**). Group 0 cells were brown adipocyte progenitors highly expressing mesenchymal marker genes including *DLK1, THY1,* and *CD34* (**Figure 4B** and **Figure S4C**). Group 1 and 3 cells gradually ceased the expression of these mesenchymal markers while began to express genes such as *FABP4, LPL, PPARG, IGFBP5, APOE,* and *PLIN2* (**Figure 4B, C** and **Figure S4C**), thus being precursor cells undergoing adipogenic differentiation. The expression of these adipocyte genes was highest in Group 2 committed brown preadipocytes (**Figure 4C** and **Figure S4C**). The transcriptional hierarchy of adipogenic stromal cells supports previous histologic findings that human iBAT is actively developing at this gestational stage (Merklin, 1974).

**Figure 4.**
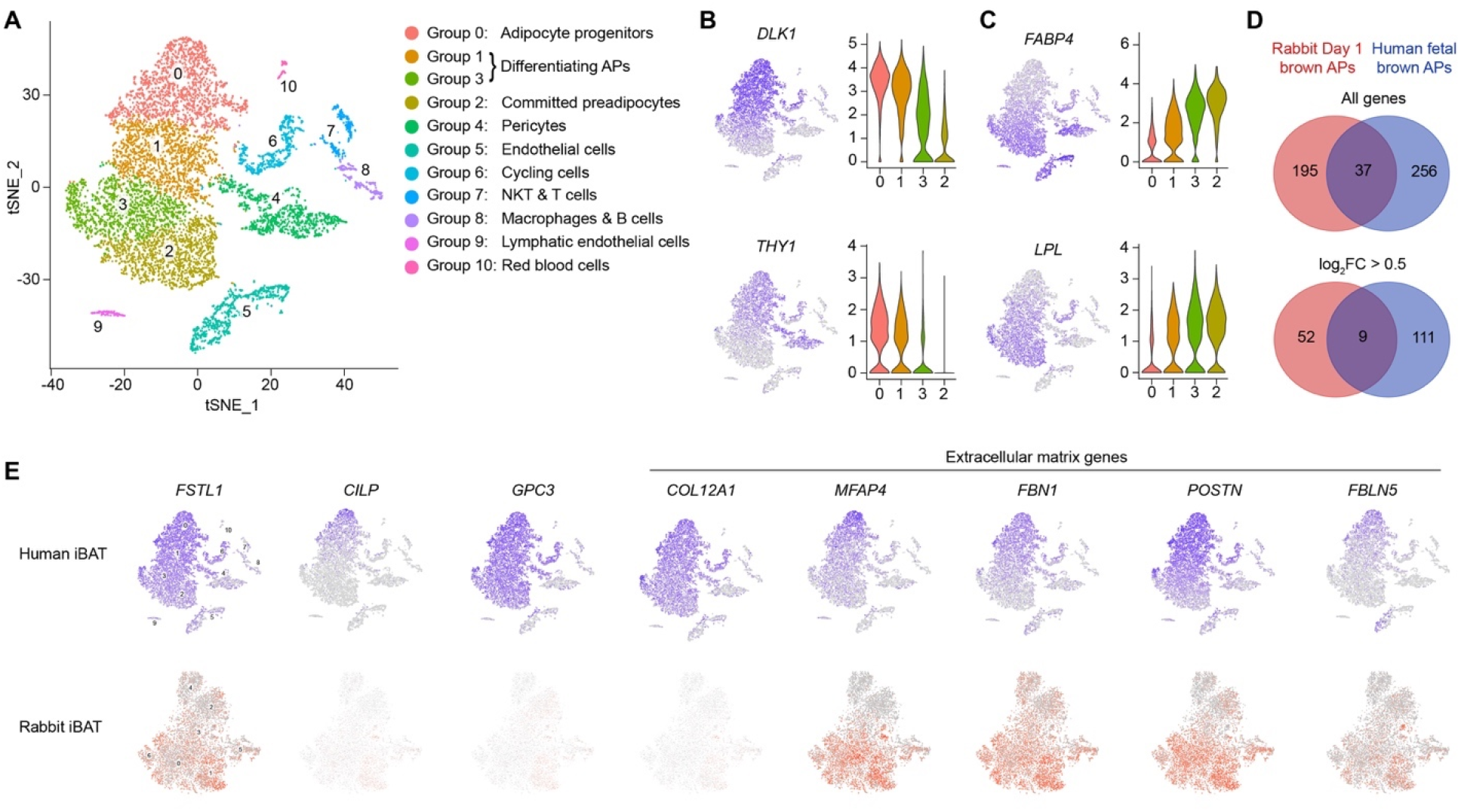
Analogy between human and rabbit brown adipocyte progenitors. (A) tSNE plot showing 11 clustered groups of SVF cells from human fetal iBAT. (B, C) tSNE and violin plots showing the expression levels and distribution of marker genes for brown adipocyte progenitors (B) and committed brown preadipocytes (C). (D) Venn diagrams of numbers of common mark genes between rabbit group #1 and human group #0 progenitors. Top, all genes. Bottom, genes with log2 fold change > 0.5. (E) tSNE plots showing the distribution of common marker genes for brown adipocyte progenitors between human (top) and rabbit (bottom).

WAT heterogeneity in mice has recently been revealed by scRNA-seq (Burl et al., 2018; Hepler et al., 2018; Schwalie et al., 2018; Cho et al., 2019; Merrick et al., 2019; Rajbhandari et al., 2019). We then profiled the expression of newly-identified white adipocyte progenitor marker genes (Rondini and Granneman, 2020) in human BAT stromal cells. *PI16* was also expressed by Group 0 cells, thus appearing as a common marker for both brown and white adipocyte progenitors (**Figure S4D**). However, *DPP4, F3 (CD142), CD55, EBF2* (Wang et al., 2014), and *ANAX3* were barely expressed nor enriched in any stromal population of cells (**Figure S4D**), suggesting that brown and white adipocyte progenitors have distinct transcriptional profiling, a manifest of their different developmental origins (Sanchez-Gurmaches et al., 2016).

To evaluate the analogy between human and rabbit brown adipocyte progenitors, we then compared upregulated marker genes of human Group 0 progenitors and rabbit Group 1 progenitors and were able to identify 37 overlapping genes (**Figure 4D** and **Table S5**). Among these, 9 genes had log2-fold-change values greater than 0.5 (**Figure 4D** and **Table S6**). These genes are *DLK1* (**Figure 3B** and **Figure 4B**), *FSTL1, CILP, GPC3, COL12A1, MFAP4, FBN1, POSTN,* and *FBLN5* (**Figure 4E**). Interestingly, 17 out the 37 overlapping genes encode extracellular proteins (**Table S7**). All the top 9 shared marker genes encode either cytokines or extracellular matrix proteins (**Table S7**), indicating the high secretory activity of brown adipocyte progenitors. Hereafter, we refer these brown adipocyte progenitors as *FSTL1*^high^ cells.

### Disappearance of FSTL1^high^ adipocyte progenitors during rabbit iBAT involution

We then sought to ask whether iBAT involution in rabbits is associated with the depletion of *FSTL1*^high^ brown adipocyte progenitors. We selected Group 1, 0, and 6 cells that were mainly comprised of iBAT adipocyte progenitors at the age of 1 day, 3 weeks, and 12 weeks, respectively (**Figure 5A**). 46 genes were identified to display higher expression in Group 1 brown adipocyte progenitors when compared to Group 0 and 6 progenitors (**Figure 5B** and **Table S8**). Out of the aforementioned 9 common markers for brown adipocyte progenitors (**Table S6**), *DLK1, FSTL1, CILP, GPC3,* and *COL12A1* reduced their expression in progenitor cells when iBAT involuted at the age of 3 and 12 weeks (**Figure 5C**).

**Figure 5.**
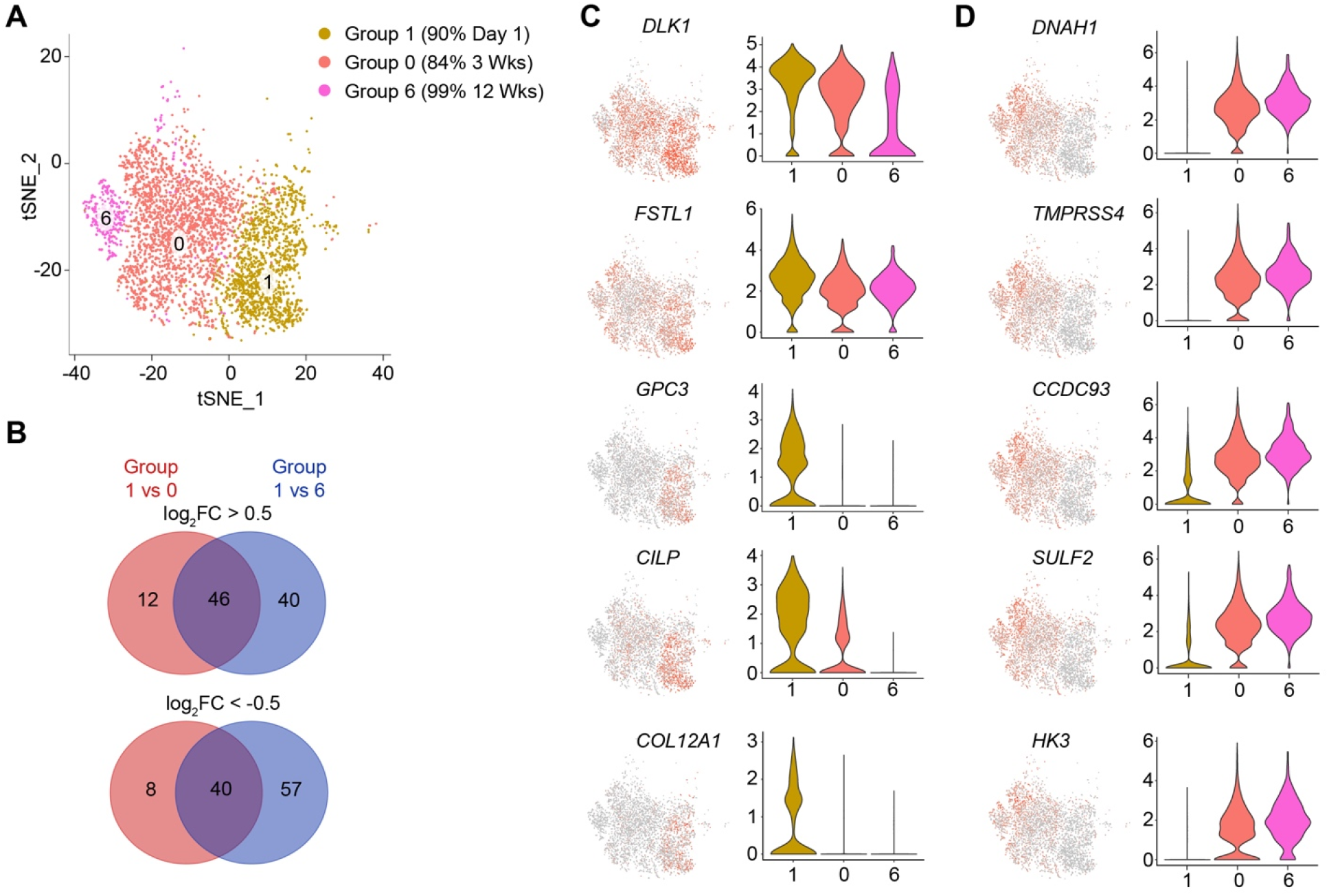
Single-cell transcriptional reprograming of adipocyte progenitors during rabbit iBAT involution. (A) tSNE plot showing group 1, 0, and 6 progenitors derived from rabbit iBAT SVF cells. (B) Venn diagrams of upregulated (top) and downregulated (bottom) marker genes in group 1 progenitors, compared to group 0 and 6 progenitors. (C, D) tSNE and violin plots showing the expression levels and distribution of marker genes for group 1 brown adipocyte progenitors (C) and involuted group 0 and 6 adipocyte progenitors (D).

40 genes, such as *DNAH1, TMPRSS4, CCDC93, SULF2,* and *HK3,* were upregulated in Group 0 and 6 progenitor cells (**Figure 5D** and **Table S9**). More importantly, these late onset genes in Group 0 and 6 cells were mostly not expressed by human BAT *FSTL1*^high^ progenitors (**Figure S5A**). On the other hand, WAT progenitor markers including *DPP4* and *PI16* started to express in Group 0 and 6 cells (**Figure S5B, C**), indicating that they became, at lease gained transcriptional properties of, white adipocyte progenitors. Thus, the transcriptional reprogramming of adipocyte progenitors and loss of *FSTL1*^high^ cells may underlie the involution process of iBAT.

### FSTL1 is a cytokine enriched in brown adipocyte progenitors

Next, we then set out to explore whether the *FSTL1* genes was functionally important for *FSTL1*^high^ brown adipocyte progenitors. The encoded glycoprotein follistatin like 1 belongs to the secreted protein acid and rich in cysteine (SPARC) family that has a role in lung development, muscle function, and cardioprotection (Mattiotti et al., 2018a). However, whether it controls iBAT development and involution has not been explored. At the RNA level, *FSTL1* was one of the highest expressed gene in total iBAT and sBAT from rabbits (**Figure S6A**). At the protein level, FSTL1 showed correlated reduction with UCP1, to minimally detectable levels in total iBAT extracts from 6- and 12-week-old rabbits (**Figure 6A**). In humans, SVF cells derived from fetal iBAT (Wang et al., 2018) had more glycosylated and un-glycosylated forms of FSTL1 (Wei et al., 2015) (**Figure 6B**), when compared to immortalized SVF cells from adult neck BAT and WAT (Xue et al., 2015). These data indicate that levels of FSTL1 are positively correlated with the brown adipogenic ability of progenitor cells in the SVF.

**Figure 6.**
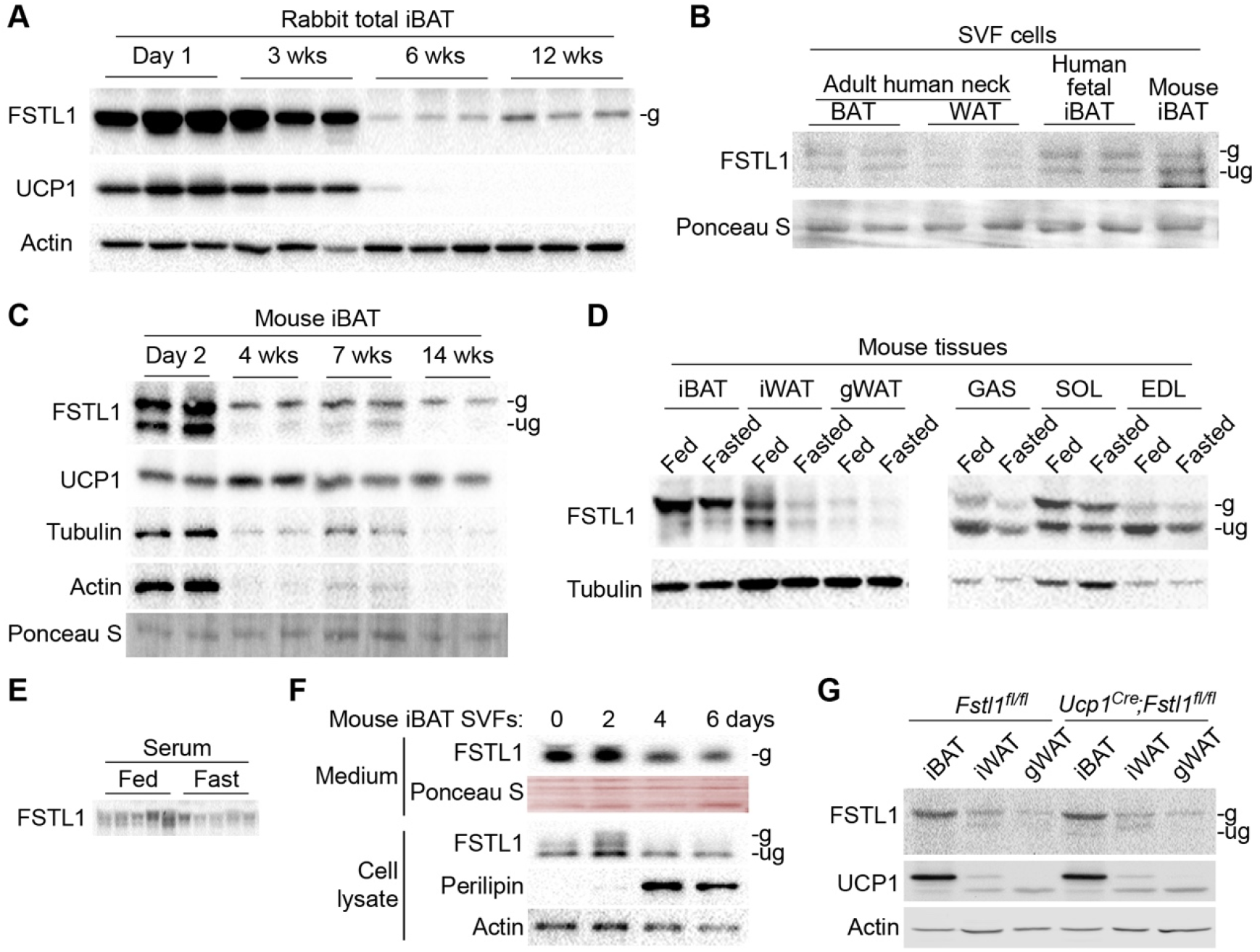
FSTL1 is a brown adipocyte progenitor-enriched factor. (A) Expression of FSTL1 and UCP1 protein in total iBAT from rabbits at indicated ages. “g” and “ug” indicate glycosylated and unglycosylated FSTL1, respectively. (B) FSTL1 protein expression in SVF cells from BAT and WAT of adult human neck, human fetal iBAT, and mouse iBAT. (C) FSTL1 and UCP1 protein expression in iBAT from mice at indicated ages. (D) FSTL1 protein expression in iBAT, inguinal and gonadal WAT, gastrocnemius (GAS), soleus (SOL), and extensor digitorum longus (EDL) muscle from mice either fed *ad libitum* or fasted for 24 h. (E) FSTL1 levels in the serum of mice either fed *ad libitum* or fasted for 24 h (n = 5). (F) FSTL1 and UCP1 protein expression in BAT and WAT depots of *Fstl1^fl/fl^* and *Ucp1^Cre^;Fstl1^fl/fl^* mice. (G) FSTL1 protein expression and secretion into medium during the adipogenic differentiation of SVF cells from mouse iBAT.

SVF cells from mouse iBAT can be readily differentiated into UCP1-positive brown adipocytes. Comparable amount of FSTL1 protein was detected in iBAT SVF cells derived from young mice and human fetus (**Figure 6B**). In newborn mice when iBAT is actively growing, FSTL1 expression was most abundant (**Figure 6C**). However, there was a notable and steady reduction in FSTL1 levels after weaning (**Figure 6C**). Compared with inguinal WAT (iWAT), gonadal WAT (gWAT) and skeletal muscle, iBAT processed highest level and mostly glycosylated form of FSTL1 (**Figure 6D**). Fasting, a condition when thermogenic activity is suppressed (Ruan et al., 2014), reduced FSTL1 levels in tissues (**Figure 6D**) and serum (**Figure 6E**) in mice. These data demonstrate that levels of glycosylated FSTL1 are associated with BAT activity.

During the adipogenesis of iBAT SVF cells in culture, FSTL1 protein expression and its secretion into the medium were initially upregulated and then drastically diminished when mature brown adipocytes were induced (**Figure 6F**). This was consistent with the finding that *FSLT1* gene expression was downregulated in committed preadipocytes of both humans and rabbits (**Figure 4E**). In support of the notion that FSTL1 is an adipocyte progenitor-enriched cytokine, deleting the *Fstl1* gene in brown/beige adipocytes or all mature adipocytes using the *Ucp1^Cre^* and the *Adipoq^Cre^,* respectively did not evidently affect levels of FSTL1 protein in iBAT, WAT or serum (**Figure 6G and Figure S6C, D**). Consistently, we did not observe any changes in UCP1 expression (**Figure 6G**), body weight, glucose metabolism, or cold-induced thermogenesis in *Ucp1^Cre^;Fstl1^fl/fl^* or *Adipoq^Cre^;Fstl1^fl/fl^* knockout mice (**Figure S6E-I**). These data indicate that mature thermogenic adipocyte-derived FSTL1, if any, is dispensable for BAT development and function.

### Loss of FSTL1 in the *Myf5*-lineage causes iBAT atrophy in mice

To assess the role of brown adipocyte progenitor-derived FSTL1 in iBAT development in vivo, we then ablated the *Fstl1* gene in the *Myf5*-expressing common progenitors for iBAT and skeletal muscle (Timmons et al., 2007). No obvious morphological changes of iBAT during embryonic development were observed in *Myf5-Fstl1* KO mice (**Figure S7A, B**). In neonatal *Myf5-Fstl1* KO mice, only residual levels of FSTL1 were detected in iBAT and the serum (**Figure 7A** and **Figure S7C**). Compared to wildtype littermates, *Myf5-Fstl1* KO mice displayed a severe defect in thermogenesis, as shown by lower UCP1 and COX4 protein expression (**Figure 7A**), reduced body temperature when separated from dams (**Figure 7B**), smaller size of iBAT (**Figure S7D**), and depleted lipid storage within iBAT (**Figure 7C**). Most *Myf5-Fstl1* KO pups died within 5 days after birth largely because of hypothermia and muscle dysfunction (suggested by the hunched posture shown in **Figure S7D**).

**Figure 7.**
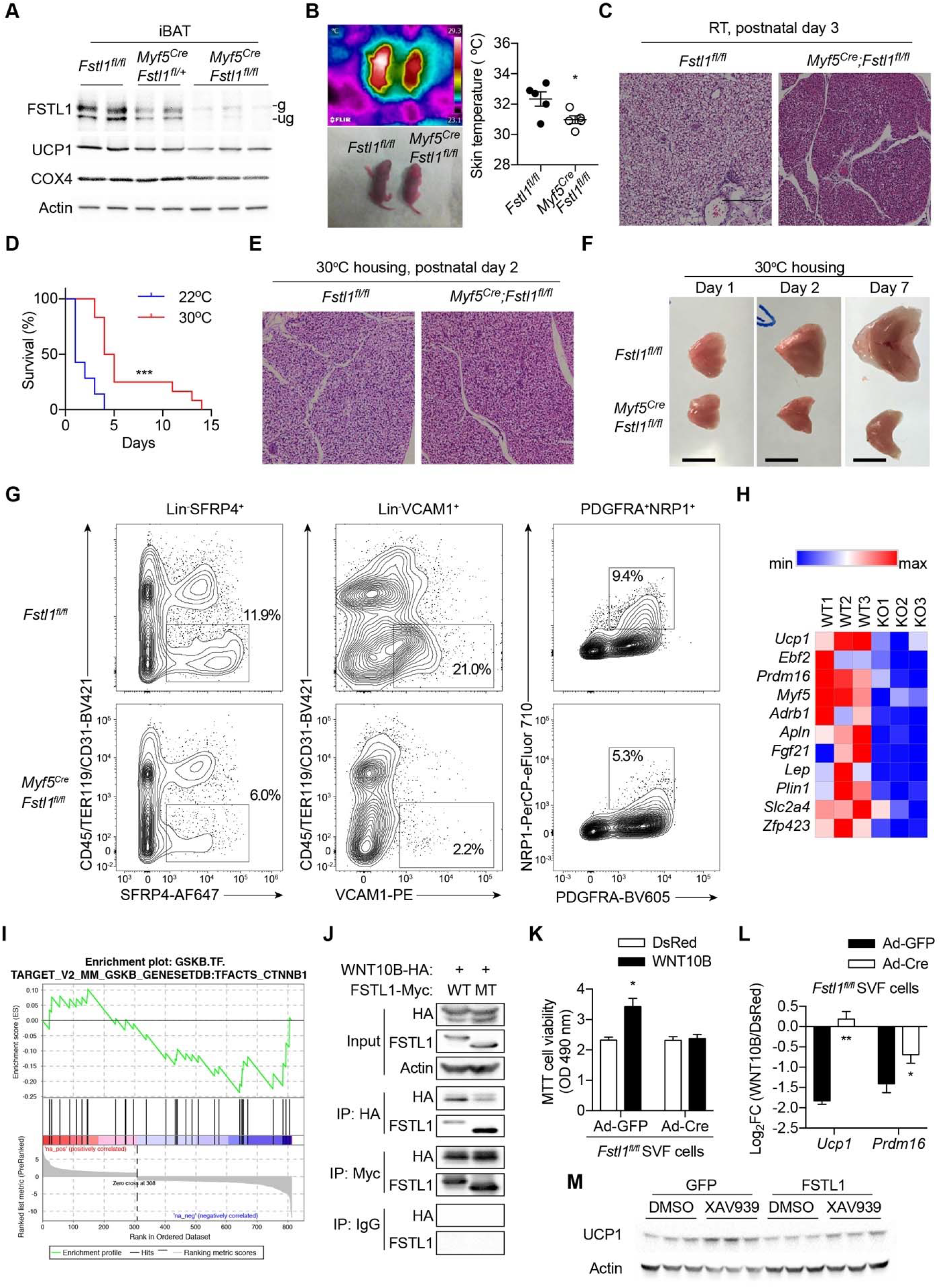
iBAT atrophy in *Myf5^Cre^;Fstl1^fl/fl^* mice. (A) Immunoblotting of FSTL1, UCP1, and COX4 in iBAT from neonatal *Fstl1^fl/fl^* control, *Myf5^Cre^;Fstl1^fl/fl^* heterozygous, and *Myf5^Cre^;Fstl1^fl/fl^* knockout mice. (B) Body surface temperature of newborn mice separated from dams at room temperature for 3 hours. Periscapular skin temperature is quantified in the right. (C) Histology of iBAT of 3-day-old control and *Myf5^Cre^;Fstl1^fl/fl^* knockout mice housed at room temperature (RT). (D) Survival curves of control and *Myf5^Cre^;Fstl1^fl/fl^* knockout mice housed at 22 or 30 °C. (E, F) Histology and morphology of iBAT of control and *Myf5^Cre^;Fstl1^fl/fl^* knockout mice housed at 30 °C. (G) Flow cytometric quantification of adipocyte progenitors (Lin^-^SFRP4^+^) and committed preadipocytes (Lin^-^SFRP4^+^ and Lin^-^PDGFRA^+^NRP1^+^) within the SVF of iBAT from control and *Myf5^Cre^;Fstl1^fl/fl^* knockout mice housed at RT. (H) Expression heatmap of selected markers of thermogenesis and adipogenesis, determined by RNA-seq. (I) Enrichment plots for β-Catenin-regulated genes from the Gene Set Enrichment Analysis (GSEA) of differentially expressed genes between control and *Myf5^Cre^;Fstl1^fl/fl^* knockout iBAT. (J) Interaction between WNT10B and wildtype (WT) and glycosylation-mutant (MT: N142/173/178Q) determined by reciprocal co-immunoprecipitation with protein lysate from overexpressed HEK 293 cells. (K, L) *Fstl1^fl/fl^* iBAT SVF cells were infected with adenoviruses expressing GFP or Cre recombinase to induce *Fstl1* gene deletion. Meantime, DsRed or WNT10B were overexpressed via lentiviral vectors. MTT proliferation assay of undifferentiated cells (K, n = 5) and qRT-PCR of *Ucp1* and *Prdm16* genes of differentiated cells (L, n = 6) were performed. Log_2_Fold change(+/- WNT10B) values are shown in L. (M) Primary iBAT SVF cells were infected with lentiviral vectors expressing GFP or FSTL1, treated with or without XAV939, and differentiated into adipocytes. UCP1 protein expression was determined by Western blotting. Data are presented as mean ± SEM.*, p < 0.05; ***, p < 0.001 by unpaired student’s t-test (B, L) Mantel-Cox test (D), and two-way ANOVA (K).

Housing mice at thermoneutrality delayed the death of *Myf5-Fstl1* KO pups (**Figure 7D**) and restored lipid content within iBAT (**Figure 7E**). Muscle defects persisted in *Myf5-Fstl1* KO mice at thermoneutral zone (data not shown), presumably causing the mortality eventually. However, iBAT defects observed in *Myf5-Fstl1* KO mice were not results of the loss of skeletal muscle-derived FSTL1, since deleting FSTL1 in skeletal myocytes using the *HSA^Cre^* line did not affect iBAT morphology, UCP1 expression, body weight, or glucose metabolism (**Figure S7E-H**). An assessment of the iBAT morphology at thermoneutrality showed that *Myf5-Fstl1* KO iBAT did not increase the mass from day 1 to 7 (**Figure 7F**). We then used novel markers identified from scRNA-seq of rabbit SVF cells (**Figure 3B, C**) for flow cytometric assays. In the SVF fraction of *Myf5-Fstl1* KO iBAT, numbers of lineage-negative SFRP4^+^ brown adipocyte progenitors and VCAM1^+^ and PDGFRA^+^NRP1^+^ committed preadipocytes were significantly reduced (**Figure 7F**). These results collectively demonstrate that FSTL1 deletion in brown adipose progenitors causes iBAT atrophy in mice.

iBAT atrophy in *Myf5-Fstl1* KO mice was not because of defects in sympathetic nerve innervation, tissue vascularization, or macrophage polarization (**Figure S7I-K**). We then performed RNA-seq of total iBAT from newborn mice and found 505 upregulated and 308 downregulated genes with more than 2-fold change in *Myf5-Fstl1* KO mice (**Table S10**). Adipogenic and thermogenic genes were enriched in those downregulated while inflammatory and immune-activating genes were upregulated (**Figure 7H** and **Figure S7L**). Gene Set Enrichment Analysis (GSEA) identified that β-Catenin-controlled genes were overrepresented in the downregulated gene set (**Figure 7I**). The WNT/β-Catenin signaling has been extensively shown to promote the self-renewal of adipocyte progenitor while inhibits adipogenic differentiation (Kang et al., 2005; Lo et al., 2016). We interrogated BioPlex 2.0, a database of protein-protein interactions profiled in human cells via affinity-purification mass spectrometry (Huttlin et al., 2017), and found that FSTL1 binds to several WNT proteins (**Table S11**). Coimmunoprecipitation of overexpressed proteins validated that WNT10B physically interacts with both FSTL1, regardless of the glycosylation status (**Figure 7J**). Moreover, WNT10B-stimulated SVF proliferation (**Figure 7M**) and -inhibited thermogenic gene expression after adipogenic differentiation (**Figure 7L**) were ablated when FSTL1 was absent (**Figure S7M**). On the other hand, inhibiting WNT signaling with XAV939 increased UCP1 protein expression in differentiated SVF cells, which was diminished by FSTL1 overexpression. These data suggest that FSTL1 may act downstream of the WNT signaling to sustain the brown adipose progenitor pool and support the postnatal expansion of iBAT.

## DISCUSSION

As an energy-consuming, endocrine organ, BAT promotes weight loss and metabolic fitness when activated or transplanted in rodents (Soler-Vazquez et al., 2018). In humans, the activity of BAT has been repeatedly reported to be inversely associated with body mass index (Betz and Enerback, 2011; Tam et al., 2012). Chronic treatment with the β3-adrenergic receptor agonist Mirabegron increases BAT activity and insulin sensitivity in young adults (O’Mara et al., 2020). Nonetheless, the factor that human BAT, including peri-scapular, peri-jugular and perirenal depots, undergoes age-related involution/atrophy raises doubts about the physiological significance of BAT, particularly in middle-aged humans. One could assume that activating residual (if any left) brown and/or beige adipocytes in mid-to late-life would have limited metabolic benefits, but we argue that preventing/delaying BAT involution or regenerating BAT via stem cell therapy and tissue engineering should have clinical relevance for counteracting obesity and its comorbidities. However, the lack of understanding of the nature and mechanisms of BAT involution impedes translational efforts towards regenerating BAT.

BAT involution is not unique to humans and has been observed in mammals such as sheep, bovines, and rabbits (Derry et al., 1972; Gemmell et al., 1972; Casteilla et al., 1989; Basse et al., 2015; Oelkrug et al., 2015). Yet, mechanistic insight into this seemly evolutionarily conserved process is largely lacking. In this study, we found that tissue whitening and identity switch of adipocytes during rabbit iBAT involution is associated with the compromised browning ability of perivascular precursors and the loss of *FSTL1*^high^ brown adipocyte progenitors. The fate of mature brown adipocytes existed in neonatal rabbits is still unclear. Whether they undergo cell death or transdifferentiation requires future fate-mapping experiments. The unilocular fat cells observed in involuted BAT are presumed to be white adipocytes, based on their morphology, gene expression, and refractoriness to cold-induced remodeling. A small proportion of these unilocular fat cells could become multilocular when activated by β3-adrenergic signaling, indicating their potential identity as beige adipocytes. However, we could not rule out the possibility that they are residual “dormant” brown adipocytes. Similarly, the fate of *FSTL1*^high^ brown adipocyte progenitors is also an open question: do they become extinct or transdifferentiate into white adipocyte progenitors? Importantly, comparable *FSTL1*^high^ progenitors can be found in human fetal iBAT. It would be of high interest in the future to determine whether the prevalence of *FSTL1*^high^ progenitors decreases as iBAT atrophies in young people. However, establishing this potential correlation or causation in humans is currently hindered by the lack of access to biological samples and the lack of simple and reliable approach to quantify *FSTL1*^high^ progenitors.

In contrast to humans and rabbits, mice have constitutively high levels of FSTL1 expressed by brown adipocyte progenitors. This could be a potential mechanism to support the life-long existence and activeness of BAT in mice to cope with the high thermogenic demand. Recently, using whole-body heterozygous *Fstl1^+/-^* mice, we showed that FSTL1 is required for optimal BAT recruitment during cold and after β3-adrenergic activation (Fang et al., 2019). The thermogenic effect of FSTL1 cannot be attributed to mature brown adipocytes, as its expression and secretion were downregulated during adipogenic differentiation and no changes in FSTL1 expression or BAT function were observed in *Ucp1^Cre^;Fstl1^fl/fl^* and *Adipoq^Cre^;Fstl1^fl/fl^* knockout mice. Consistently with the enrichment of FSTL1 in brown adipocyte progenitors, only deleting FSTL1 in *Myf5^+^* cells diminished brown adipose progenitors and caused iBAT atrophy and thermogenic defects in mice. Skeletal muscle cells are also derived from the *Myf5^+^* lineage. Consequently, muscle defects were apparent in *Myf5^Cre^;Fstl1^fl/fl^* knockout mice, which contributed to the neonatal mortality. However, we believe that the role of FSTL1 in BAT involution is tissue-autonomous, because BAT development and function were intact in *HSA^Cre^;Fstl1^fl/fl^* knockout mice, even though serum FSTL1 was dropped to a level similar to that in *Myf5^Cre^;Fstl1^fl/fl^* knockout mice. We also argue that the contribution of muscle satellite cells to BAT maintenance should be minimal, as these cells are rare, quiescent, and acting locally (Yin et al., 2013). Nonetheless, validating the cell-intrinsic function of FSTL1 requires selective gene targeting in brown adipocyte progenitors, but genetic tools are still lacking. Besides BAT, FSTL1 is also expressed in WAT. It is transiently induced during the adipogenic differentiation of 3T3-L1 cells and then rapidly downregulated to background level (Wu et al., 2010; Fang et al., 2019). Intriguingly, either knocking down *Fstl1* or treatment with high levels of recombinant FSTL1 interfere the 3T3-L1 differentiation (Prieto-Echague et al., 2017). Our unpublished results and publicly available scRNA-seq data both suggest FSTL1 as an enriched factor for mesenchymal cells in various tissues (Tabula Muris et al., 2018). Therefore, we speculate that FSTL1 is not a marker distinguishing brown versus white adipocyte progenitors, but rather an important factor maintaining the quiescence and identity of mesenchymal stem/progenitor cells.

Our scRNA-seq data also demonstrate that genes encoding secreted proteins are clustered in *FSTL1*^high^ brown adipocyte progenitors in both humans and rabbits. These genes could be preferentially identified simply because they are highly expressed, but also suggesting the high secretory activity of *FSTL1*^high^ progenitors. The study of brown adipokines (batokines) has been mainly focused on autocrine, paracrine, and endocrine factors from mature brown adipocytes (Villarroya et al., 2017), yet the contribution of perivascular precursors has so far largely neglected. Among these common extracellular factors expressed in *FSTL1*^high^ cells, DLK1 (also known as PREF1) is a well-established marker for mouse adipocyte progenitors that also inhibits adipogenesis (Smas and Sul, 1993; Hudak et al., 2014). FSLT1 displays comparable expression patterns to DLK1, and thus may have an analogous function in adipogenesis. FSTL1 is involved in multiple signaling pathways, and many of its biological functions have been attributed to its inhibition to BMP signaling by competitively binding to BMP4 (Geng et al., 2011; Mattiotti et al., 2018b). However, BMP signaling has been widely recognized to promote the commitment of mesenchymal stem cells into the adipogenic lineage (Tang and Lane, 2012; Lee, 2018). It would be counterintuitive if FSTL1 in BAT inhibits BMP signaling, which would predict increased adipogenic commitment and differentiation in *Myf5^Cre^;Fstl1^fl/fl^* knockout mice. Nonetheless, the BMP4-antagonistic effect of FSTL1 might determine the adipocyte identity in BAT, since BMP4 can shift brown adipocytes to white-like fat cells (Modica et al., 2016). In our study, RNA-seq of *Myf5^Cre^;Fstl1^fl/fl^* iBAT and subsequent bioinformatic analysis pointed to reduced WNT/β-Catenin activity, which has been extensively shown to promote the proliferation and potency maintenance of adipose progenitor cells (Prestwich and Macdougald, 2007; Chen and Wang, 2018). Indeed, we found that FSTL1 physically interacts with WNT10B in vitro and acts downstream of WNT10B to balance the proliferation and the differentiation of SVF cells. Nonetheless, future studies are required to determine the molecular underpinnings of FSTL1-WNT interplay.

Studies in rodents have proposed several mechanisms for age-associated functional decline of brown and beige fat. Externally, impairment of sympathetic input, altered endocrine control, and tissue inflammation during aging contribute to thermogenic dysfunction (Graja et al., 2019; Zoico et al., 2019). Adipocyte-intrinsic mechanisms include reduced UCP1 expression and activity (Horan et al., 1988; McDonald et al., 1988; Florez-Duquet et al., 1998), cellular senescence (Berry et al., 2017), and mitochondrial lipoylation (Tajima et al., 2019b). However, it is unclear whether these deteriorations that normally happen in late life play a part in iBAT involution during early life of humans and rabbits. Our study is a further step towards the understanding of this multifactorial process. Moreover, cervical and supraclavicular BAT in adult humans also goes through age-dependent involution. Whether *FSTL1*^high^ progenitors also exist in these BAT depots and whether their disappearance also drives the involution of these BAT depots await future investigation.

In summary, we propose that rabbits can serve as an ideal model, alternatively to rodents, for the study of BAT involution. We define brown adipocyte progenitors as *FSTL1*^high^ cells in iBAT of both humans and rabbits. Age-dependent transcriptional and functional remodeling of iBAT is associated with the depletion of *FSTL1*^high^ progenitors in rabbits. Loss of FSTL1 in brown adipocyte progenitors attenuates WNT signaling and drives iBAT atrophy in mice. Our data could potentially inform future development of preventive and regenerative medicine in obesity and comorbidities.

## ACKNOWLEDGEMENT

We thank Dr. Maria Razzoli and Dr. Pilar Ariza Guzman for EchoMRI and CLAMS analyses, Dr. Xiang Gao, Dr. Xu Zhang and Dr. David Broide for providing the *Fstl1-floxed* mice, Dr. Yu-Hua Tseng for providing immortalized human brown preadipocytes, and Dr. Chenbo Ji for providing embryonic human brown preadipocytes. We thank the UMN Research Animal Resources for assisting in rabbit husbandry and procedures. This work was supported by National Natural and Science Foundation of China (31500944), National Key R&D Program of China (2017YFD0500505), Natural Science Foundation of Jiangsu Province (BK20150687), and China Scholarship Council postdoctoral fellowship (201606855010) to Z. H.; Natural Science Foundation of Jiangsu Province (BK20170147) to Z. Z.; American Diabetes Association (18-IBS-167) and NIH/NIAID (R01 AI139420 and R21 AI140109) to H.-B.R. H.N. and S.S. were supported by NIH grants UL1 TR002494 and R01 AI148669.

## AUTHOR CONTRIBUTIONS

Z.H. and Z.Z. designed and performed experiments, analyzed data, and wrote the manuscript. R.H. and K. R. assisted in mouse colony management and experiments. P. H. collected human fetal iBAT. Z.S and D.C. performed longitudinal transcriptomic analyses of bulk RNA-seq of SVF cells. A.B. helped with metabolic phenotyping. Y.G. helped with the generation of *Fstl1* KO mice. W. Z. obtained funding. H.N. and S.S. analyzed rabbit scRNA-seq data. H.-B.R. conceived the project, designed experiments, analyzed data, and wrote the manuscript.

