## Supplemental Methods for "Loss of FSTL1-expressing adipocyte progenitors drives the age-related involution of brown adipose tissue"

**KEY RESOURCES TABLE**

| **REAGENT or RESOURCE** | **SOURCE** | **IDENTIFIER** |
| --- | --- | --- |
| Antibodies | | |
| Anti-UCP1 (Dilution 1:5000 WB& 1:5000 IHC) | Abcam | Cat# ab209483 |
| Anti-Tubulin (Dilution 1:2000 WB) | Proteintech | Cat# 66031-1-1g |
| Anti-FSTL1 (Dilution 1:500 WB) | R&D Systems | Cat# AF1738  LOT# JRB0115071 |
| Anti-Actin (Dilution 1:5000 WB) | Sigma-Aldrich | Cat# 5441 |
| Anti-Perilipin (Dilution 1:1000 WB) | Cell Signaling Technology | Cat# 9349  LOT# 4 |
| Anti-GAPDH (Dilution 1:2000 WB) | Bioworld | Cat# MB001  LOT# XS2018012026 |
| Anti-COX4 (Dilution 1:1000 WB) | Proteintech | Cat# 11242-1-AP  LOT# 00042713 |
| Anti-β-Catenin (Dilution 1:1000 WB) | Cell Signaling Technology | Cat# 8480  LOT# 5 |
| Anti-HA (2 μg IP) | Proteintech | Cat# 51064-2-AP  LOT# 00061770 |
| Anti-HA (Dilution 1:1000 WB) | Roche | Cat# 11583816001  LOT# 37081400 |
| Anti-Myc (2 μg IP) | Proteintech | Cat# 16286-1-AP  LOT# 00066621 |
| Anti-TH (Dilution 1:1000 WB) | Cell Signaling Technology | Cat# 13106  LOT# 2 |
| Anti-CD31 (Dilution 1:100 IF) | Bioworld | Cat# BS1574 |
| Anti-CD45-BV421 (Dilution 1:100 FACS) | BD Bioscience | Cat# 564279  LOT# 9016570 |
| Anti-TER119-BV421 (Dilution 1:100 FACS) | BioLegend | Cat# 116234  LOT# B260380 |
| Anti-CD31-BV421 (Dilution 1:100 FACS) | BioLegend | Cat# 102424  LOT# B265877 |
| Anti-SFRP4 (Dilution 1:100 FACS) | Abcam | Cat# ab154167  LOT# GR3182439-6 |
| Anti-VCAM1-PE (Dilution 1:100 FACS) | BioLegend | Cat# 105713  LOT# B265630 |
| Anti-NRP1-PerCP-eFluor710 (Dilution 1:100 FACS) | Thermo Fisher Scientific | Cat# 46-3041-80  LOT# 1949804 |
| Anti-PDGFRA-BV605 (Dilution 1:100 FACS) | BioLegend | Cat# 135916  LOT# B260842 |
| Anti-Mouse IgG-HRP (Dilution 1:10000 WB) | Cell Signaling Technology | Cat# 7076 |
| Anti-Rabbit IgG-HRP (Dilution 1:10000 WB) | Cell Signaling Technology | Cat# 7074 |
| Anti-Goat IgG-HRP (Dilution 1:10000 WB) | Santa Cruz Biotechnology | Cat# SC2020 |
| Anti-Rabbit IgG-AF647 (Dilution 1:200 FACS) | BioLegend | Cat# 406414  LOT# B281520 |
| Anti-Rabbit IgG-AF488 (Dilution 1:1000 IF) | Thermo Fisher Scientific | Cat# R37118 |
| Normal Rabbit IgG (2 μg IP) | Santa Cruz Biotechnology | Cat# SC-2027  LOT# A2412 |
| Bacterial and Virus Strains | | |
| Adenovirus-EGFP | Ruan et al., 2012 | N/A |
| Adenovirus-Cre | Ruan et al., 2012 | N/A |
| Chemicals, Peptides, and Recombinant Proteins | | |
| DMEM | Corning | Cat#10-017-CV |
| DMEM/F12 | Corning | Cat#10-090-CV |
| Adipocyte Medium | Sciencell | Cat#7201 |
| FBS | GenClone | Cat#25-514 |
| PBS | Corning | Cat#20-040-CV |
| HBSS | Thermo Fisher Scientific | Cat#14175-095 |
| HEPAS | Thermo Fisher Scientific | Cat#15630-080 |
| Penicillin-Streptomycin (Pen-Strep) | Thermo Fisher Scientific | Cat#15140122 |
| 70um Filter | Fisher Brand | Cat#22363548 |
| Collagenase I | Worthington | Cat#LS004196 |
| Liver Digest Medium | Thermo Fisher Scientific | Cat# 17703-034 |
| 40um Filter | Fisher Brand | Cat#22363549 |
| ACK Buffer | Quality Biological | Cat#118-156-101 |
| Ciprofloxacin HCl | Sigma-Aldrich | Cat#PHR1044 |
| Blasticidin | InvivoGen | Cat#ant-bl |
| Primocin | InvivoGen | Cat#ant-pm-1 |
| MTT (3-(4,5-Dimethylthiazol-2-yl)-2,5-Diphenyltetrazolium Bromide) | Sigma-Aldrich | Cat#M2128 |
| PEI Max | Polysciences | Cat# 24765-1 |
| Insulin (Novolin R^®^) | Novo Nordisk | Cat# 0169-1833-11 |
| T3 (3,3’,5-Triiodo-L-thyronine sodium salt) | Sigma-Aldrich | Cat#T6397 |
| IBMX (3-Isobutyl-1-methylxanthine) | Sigma-Aldrich | Cat#I5879 |
| Rosiglitazone | Sigma-Aldrich | Cat#R2408 |
| Dexamethasone | Sigma-Aldrich | Cat#D4902 |
| Indomethacin | Sigma-Aldrich | Cat#I7378 |
| Biotin | Sigma-Aldrich | Cat# B4639 |
| Transferrin | Sigma-Aldrich | Cat# T8158 |
| D-Pantothenic Acid | RPI | Cat#P55650 |
| Primocin | InvivoGen | Cat#ant-pm-1 |
| 2-Deoxy-D-Glucose (2-DG) | Sigma-Aldrich | Cat#D6134 |
| Mirabegron | XA BC-Biotech | N/A |
| CL316,243 | Tocris | Cat# 1499 |
| TRIzol | Thermo Fisher Scientific | Cat# 15596026 |
| Matrigel | Coring | Cat#356237 |
| Isoflurane | Piramal | Cat#66704-013-025 |
| DMSO | Sigma-Aldrich | Cat#D4540 |
| KolliphorEL | Sigma-Aldrich | Cat#C5135 |
| Protein A/G | Santa Cruz Biotechnology | Cat#SC-2027 |
| Formalin solution | Sigma-Aldrich | Cat#HT501128 |
| VECTASHIELD® Antifade Mounting Medium with DAPI | Vector | Cat#H-1200 |
| Target Retrieval Solution, Citrate pH 6.1 | Agilent | Cat#S1699 |
| Critical Commercial Assays | | |
| Dead cell removal kit | Miltenyi Biotec | Cat#130-090-101 |
| Fixable Viability Dye eFluor™ 780 | eBioscience | 65-0865-14 |
| PureLink™ RNA Mini Kit | Thermo Fisher Scientific | Cat#12183018A |
| iScript™ cDNA Synthesis Kit | Bio-Rad | Cat#1708891 |
| iTaq™ Universal SYBR^®^ Green Supermix | Bio-Rad | Cat#172-5125 |
| Histostain®-Plus 3rd Gen IHC Detection Kit | Thermo Fisher Scientific | Cat#85-9073 |
| Deposited Data | | |
| RNA-Seq of iBAT from Myf5-Fslt1 KO mice | This paper | GSE148888 |
| Bluk and scRNA-Seq of rabbit BAT | This paper | GSE148891 |
| Experimental Models: Cell Lines | | |
| hTERT A41hBAT-SVF | ATCC | CRL-3385 |
| hTERT A41hWAT-SVF | ATCC | CRL-3386 |
| Human fetal BAT SVF | Xue et al., 2015 | N/A |
| Mouse iBAT SVF | This paper | N/A |
| Rabbit iBAT SVF | This paper | N/A |
| 293FT | Thermo Fisher Scientific | Cat#R70007 |
| Experimental Models: Organisms/Strains | | |
| *Fstl1^fl/fl^*: B6-Fstl1^f3,4/+^/Nju | National Resource Center for Mutant Mice of China | Stock No: B000172 |
| *Ucp1-Cre*: B6.FVB-Tg(Ucp1-cre)1Evdr/J | The Jackson Laboratory | Stock No: 024670 |
| *Myf5-Cre*: B6.129S4-Myf5^tm3(cre)Sor^/J | The Jackson Laboratory | Stock No: 007893 |
| *Adipoq-Cre*: B6;FVB-Tg(Adipoq-cre)1Evdr/J | The Jackson Laboratory | Stock No: 010803 |
| *HSA-Cre*: B6.Cg-Tg(ACTA1-cre)79Jme/J | The Jackson Laboratory | Stock No: 006149 |
| NCG: NOD-Prkdc^em26Cd52^Il2rg^em26Cd22^/NjuCrl | Charles River | 572 |
| New Zealand White Rabbits | Bakkom Rabbitry | N/A |
| Oligonucleotides | | |
| Table S12 | This paper | N/A |
| Recombinant DNA | | |
| Plasmid: pLenti-SV40T-P2a-Blasticidin | This paper | N/A |
| Plasmid: pLXSN2-Wnt10b-HA (NM_011718.2) | Kang et al., 2005 | N/A |
| Plasmid: pLVX-FSTL1-Myc WT(NM_008047.5) | This paper | N/A |
| Plasmid: pLVX-FSTL1-Myc MT(N142Q;N173Q;N178Q) | This paper | N/A |
| Plasmid: pSPAX2 | Addgene | #12260 |
| Plasmid: pMD2G | Addgene | #12259 |
| Plasmid: pLVX-Dsred-MonoN1 | Clontech | Cat#632152 |
| Plasmid: pLVX-Wnt10b (NM_011718.2) | This paper | N/A |
| Software and Algorithms | | |
| FLIR Tools | 5.13 | https://www.flir.com/products/flir-tools/ |
| Excel | v16.16.21 | https://www.microsoft.com |
| Graphpad | v7.0 | https://www.graphpad.com |
| FlowJo | v10 | https://www.flowjo.com |
| SOAPnuke | v1.5.2 | https://github.com/BGI-flexlab/SOAPnuke |
| HISAT2 | v2.0.4 | http://daehwankimlab.github.io/hisat2/ |
| Bowtie2 | v2.2.5 | http://bowtie-bio.sourceforge.net/bowtie2 |
| RSEM | v1.2.12 | http://deweylab.biostat.wisc.edu/RSEM |
| DEseq2 | v1.26.0 | https://bioconductor.org/packages/release/bioc/html/DESeq2.html |
| Cell Ranger | v3.0.1  v3.1.0 | https://www.10xgenomics.com |
| Seurat | v2.3  v3.1.5 | https://satijalab.org/seurat/ |
| Other | | |

**Lead Contact and Materials Availability**

Lead author: Hai-Bin Ruan

**Data and Code Availability**

RNA-seq data reported in this paper have been deposited in the Gene Expression Omnibus with the accession number GSE148888 and GSE148891.

**EXPERIMENTAL MODEL AND SUBJECT DETAILS**

**Animals**

All animal experiments were approved by the institutional animal care and use committee of the University of Minnesota. All the mice group-housed in light/dark cycle- (6am-8pm light), temperature- (21.5 ± 1.5 ^o^C), and humidity-controlled (30-70%) room, and free to access water and regular chow (Teklad #2018) unless otherwise indicated. All mice were maintained on a C57BL6 background. *Fstl1^fl/fl^* mice (#B000172) were a kind gift from Dr. Xu Zhang. *UCP1-Cre* (#024670), *Myf5-Cre* (#007893), *HAS-Cre* (#006149), and *Adipoq-Cre* (#010803) were purchased from the Jackson laboratory. For *UCP1^Cre^:Fstl1*, *Adipoq^Cre^:Fstl1*, *HSA^Cre^:Fstl1* animals, female *Fstl1^fl/fl^* mice were bred with male *Fstl1^fl/fl^:Cre* mice to get age- and gender-matched pairs for experiments. For *Myf5^Cre^:Fstl1* animals, *Fstl1^fl/fl^* females were inbreeded with *Fstl1^fl/+^:Myf5-Cre* males. 6-week and 12-week old New Zealand White rabbits were purchased from Bakkom Rabbitry and individually housed in a temperature- (19 ± 1^o^C) and humidity-controlled (30-70%) room with *ad libitμm* access to food (Envigo, #2031) and water. To obtain newborn and 3-week-old rabbit kits, pregnant does were purchased and individually housed as indicated above, fed with a different food (Envigo, #2030).

**Human subjects**

Human interscapular BAT tissue was obtained at Nanjing Maternity and Child Health Care Hospital (Nanjing, China) from a spontaneously aborted fetus with congenital heart disease at the gestational age of 24 weeks. Written informed consent was signed by parents. This study was approved by the Medical Ethics Committee of Nanjing Maternity and Child Health Care Hospital (Permit number: [2019] KY-081) and complied with the Population and Family Planning Law of China.

**METHOD DETAILS**

**Human cell culture**

Human hTERT A41hBAT-SVF and hTERT A41hWAT-SVF cells were maintained in DMEM containing 10% FBS and 1% Pen-Strep. Primary human fetal iBAT SVF cells were maintained in Adipocyte Medium and experiments were on cells with less than 5 passages. HEK 293FT cells were maintained in DMEM containing 10% FBS and 1% Pen-Strep.

**Mouse SVF isolation, culture, immortalization, and differentiation**

iBAT depots from six 1-month-old mice were collected, pooled and minced in 10 ml of digestion buffer (DMEM/F12 with 1mg/ml Collagenase I, 1% FBS, 1% HEPES, and 1% Pen-Strep). After shaking in 37 ^o^C at 100 rpm for 45 min, digested tissues were filtered thought 70-μm strainers and centrifuged at 1500 rpm for 3 min. The pellets were resuspended in ACK buffer and put on ice for 5 min to remove red blood cells. The ACK buffer was neutralized with 5 ml of DMEM/F12 plus 10% FBS and removed after a 1500 rpm centrifugation for 3 min. SVF cells were seeded in a 60-mm dish with DMEM/F12 containing 20% FBS, 1% Pen-Strep, and 10 μg/ml Ciprofloxacin. After 12 h, adherent SVF cells were infected with lentivirus expressing CMV-SV40T-P2a-Blasticidin. When confluent, infected SVF cells were passed into a 10-cm dish, cultured with DMEM/F12 (10% FBS, 1% Pen-Strep, 10 μg/ml Blasticidin), and labeled as passage 1 (P1). After an additional subculture, the immortalized P2 SVF cells were harvested into 10 cryogenic vials from a 15-cm dish for cryopreservation in liquid nitrogen. Cells with less than 5 passages were used.

For adipogenic differentiation, confluent cells were induced with DMEM/F12 containing 10% FBS, 1x Pen-Strep, 20 nM insulin, 1 nM T3, 0.5 mM IBMX, 5 μM dexamethasone, and 125 μM indomethacin. Two days later, cells were maintained in DMEM/F12 containing 10 %FBS, 1x Pen-Strep, 20 nM insulin, and 1 nM T3. Medium was changed every other day until lipid droplets appeared.

**Rabbit SVF isolation, culture, and differentiation**

Rabbit BAT SVF cells were isolated following the aforementioned mouse protocol, without immortalization and Pen-Strep was changed to Primocin. For adipogenic differentiation, cells were cultured in DMEM/F12 with 10% FBS and 1X Primocin until confluent, induced with DMEM/F12 containing 2% FBS, 1X Primocin, 3.5 μg/ml insulin, 1 nM T3, 0.5 mM IBMX, 5 μM dexamethasone, 125 μM indomethacin, 33 μM biotin, 10 μg/ml transferrin, 17 μM pantothenate, 1 μM rosiglitazone, 50 μM 2-DG. The medium was changed once 2 days later. Afterward, cells were maintained in DMEM/F12 with 2% FBS, 1X Primocin, 3.5 μg/ml insulin, 1 nM T3, 1 μM rosiglitazone, changed every other day until lipid droplets appeared.

**Matrigel implantation**

Rabbit SVF cells were cultured in DMEM/F12 with 10% FBS and 1X Primocin until being confluent. Cells were then washed with PBS and detached with trypsin-EDTA at 37 ^o^C for 5 min. After neutralized with culture medium, 10^7^ cells were pelleted at 300 g for 3 min, resuspended in 1 ml of Matrigel, kept on ice, and injected subcutaneously into the back of a 6-week-old NCG mouse with an insulin syringe. Slowly retrieve the needle during injection to obtain a flat implantation.

**β3 adrenergic receptor agonism**

Mirabegron stock was prepared in DMSO at 175 mg/ml. 900 μl mirabegron stock was further dissolved in 44.1 ml of 5% Kolliphor EL solution and filtered through a 0.22-μm filter. Rabbits received daily intraperitoneal injections at 3.5 mg/kg body weight for 14 days. CL316,243 stock was prepared in DMSO at 4.8823 mg/ml. 100 μl stock was further dissolved in 2341.6 μl of 0.9% NaCl solution and sterilized with a 0.22-μm filter. Mice received daily intraperitoneal injections at 1 mg/kg body weight for 7 days.

**Glucose and insulin tolerant tests**

For glucose tolerance test, 16 h-fasted mice were intraperitoneally injected with glucose (20% in saline, 1.5 g/kg body weight). Blood glucose from tail-vein blood collected at the designated times was measured using a Bayer Contour Glucometer. For insulin tolerant test, mice were fasted for 6 h, and blood glucose was recorded after Insulin (100 IU/μl, 0.75 IU/kg body weight) was injected intraperitoneally.

**Body temperature**

To measure skin temperature of mice, pups were separated from dams, briefly placed in a 30 ^o^C incubator, followed by the mild cold challenge at 22 ^o^C for 3 h. Pictures were taken with a FLIR-C2 thermal camera and peri-scapular skin temperature was measured with the FLIR tool. To measure rectal temperature during cold challenge, individually housed mice were placed in their home cages at 4 °C, with free access to food and water. A fully lubricated rectal probe (Physitemp) was used to record core body temperature at indicated time points.

**Histology**

Adipose tissues were fixed in formalin solution at 4 ^o^C for 24 h. Tissue embedding, sectioning, and hematoxylin and eosin staining were performed at the Comparative Pathology Shared Resource of the University of Minnesota. For immunostaining, antigen retrieval was performed in Citric buffer using a 2100 Retriever (Aptum Biologics). After incubation with blocking buffer (3% BSA in PBS) for 1 h, sections were immersed with primary antibody in blocking buffer overnight at 4^o^C. For UCP1 immunohistochemistry, slides were labeled with Histostain^®^-Plus 3rd Gen IHC Detection Kit next day. For CD31 immunofluorescence, PBS-washed slides were incubated with an anti-Rabbit IgG-AF488 secondary antibody at room temperature for 1 h, and then mounted with VECTASHIELD® Antifade Mounting Medium with DAPI after three times of PBS wash. A Nikon system was used for imaging.

**Real-time RT-PCR**

RNA was isolated with Trizol and reverse transcribed into cDNA with the iScript™ cDNA Synthesis Kit. Real-time RT-PCR was performed using iTaq™ Universal SYBR^®^ Green Supermix and gene-specific primers on a Bio-Rad C1000 Thermal Cycler.

**MTT assay**

1000 cells were initially seeded into one well on a 96-well plate with 100 μl DMEM/F12 plus 2% FBS and MTT assays were performed at indicated time points. Briefly, 10 μl of MTT (5 μg/μl) was added into each well. After incubation at 37 ^o^C for 3 h, liquid was removed, and 100 μl DMSO was added to dissolve the MTT formazan. The absorbance was read at 590 nm.

**Lentivirus packaging**

HEK 293FT cells were seeded into 60-mm dishes a night before the experiment to get 70% confluence at transfection. For a 60-mm dish, the culture medium was replaced with 4 ml of fresh DMEM containing 2% FBS 1 h before transfection. 2 μg transfer plasmid, 1.5 μg pSPAX2, 0.5 μg pMD2g were mixed with 16 μl PEI Max solution (1 mg/ml) in 0.5 ml of 0.9% NaCl and set in room temperature for 20 min to form the transfection complex. After being incubated with cells for 6 h, the transfection complex was replaced with 5 ml DMEM plus 10% FBS. The medium containing lentivirus was collected every 24 h for 3 days, filtered thought a 0.45-μm PVDF filter, and kept at -80 ^o^C until use.

**Co-immunoprecipitation**

293FT cells cultured in 6-mm dishes were transfected with 2 μg pLXSN2-WNT10b-HA and 2 μg pLVX-FSTL1-Myc (WT or MT). 48 h later, protein was extracted with 1400 μl NP-40 lysis buffer. 2 μg antibody (Myc, HA, or Normal rabbit IgG) and 15 μl Protein A/G beads were added into 400 μl protein lysate and incubated overnight at 4 ^o^C. Protein A/G beads were washed with TBS for 3 times, boiled with 2x Laemmli buffer, and subjected to Western blotting.

**Flow cytometry**

SVF cells from mouse iBAT were isolated as described above. Each iBAT depot was lysed with 3 ml Collagenase I buffer in individual tubes and fileted with a 40-μm filter. SVF cells were stained with primary antibody at 1:100 dilution on ice for 30 min. For SFRP4, SVF cells were then stained with an anti-rabbit IgG-AF647 secondary antibody (1:200 dilution) for another 30 min. Fixable Viability Dye was used to exclude dead cells as instructed by the manufacturer. Flow cytometry was performed on an LSR Fortessa H0081 or X20 and analyzed with FlowJo.

**Bulk RNA-seq and longitudinal analyses**

RNA of total BAT depots or SVF cells from rabbits or mice was isolated with PureLink™ RNA Mini Kit, following the manufacturer’s instruction. RNA quality was determined by an Agilent 2100 Bioanalyzer. Library was prepared at BGI and sequencing was performed using the BGISEQ-500 platform. Reads were filtered with SOAPnuke and mapped to genome with HISAT2. Clean reads were mapped to the mm10 reference with Bowtie2 and gene expression levels were determined with RSEM. Differentially expressed genes were detected using DEseq2 with the following parameters: fold change >= 2.00 and adjusted p-value <= 0.05. The longitudinal analysis was implemented using the log10-transformed FPKM (Fragments Per Kilobase Million) and considering Tukey’s method with null hypotheses for each longitudinal pattern. For example, if simultaneous hypothesis testing of two null hypotheses “D1 – W3 ≥ 0” and “W3 – W6 ≥ 0” are rejected, they are determined as the genes with “continued up-regulation” (D1 < W3 < W6). Pathway analyses and upstream regulator analyses were performed using Ingenuity Pathway Analysis (Qiagen).

**Rabbit SVF scRNA-seq and analyses**

Rabbit iBAT depots were dissected at indicated ages and minced with sharp scissors in Liver Digest Medium (Thermo Fisher, 2 ml/g tissue). After shaking in 37 ^o^C at 100 rpm for 30 min, the digestion mix was centrifuged at 300 rpm for 10 min at 4^o^C. The pellet was resuspended in 5 ml ACK buffer and placed on ice for 5 min. After being neutralized with 5 ml of DMEM/F12 plus 10% FBS, SVF cells were filtered thought a 40-μm strainer, and pelleted by centrifugation at 300 rpm for 10 min at 4 ^o^C. SVF cells were then treated with a Dead cell removal kit (Miltenyi Biotec) and immediately subjected to single-cell library preparing with the 10X Genomics platform and a Chromium Single Cell 5’ Reagent Kit at the University of Minnesota Genomics Center. Sequencing of the library (paired-end 100bp) was performed on an Illumina NovaSeq 6000 instrument. The raw data were processed using the 10x Genomics Cell Ranger package (version 3.0.1). The rabbit transcriptome was generated by filtering genome assembly (Oryctolagus_cuniculus.OryCun2.0.dna.toplevel.fa) for protein-coding genes defined in GTF file (Oryctolagus_cuniculus.OryCun2.0.94.gtf). The data matrix generated was subsequently quality accessed and analyzed with R package Seurat (version 2.3). The default parameters were used for data scaling, normalization, PCA and clustering analysis. The cutoff threshold was 1000 genes per cell and 10% of mitochondria genes.

**Human SVF scRNA-seq and analyses**

Human fetal iBAT was digested with Collagenase I buffer (2 ml/g tissue). Collagenase I buffer was freshly prepared (44.5 ml DMEM/F12, 2 ml of 20 mg/ml collagenase I in HBSS, 500 μl HEPES, 500 μl FBS, 500 μl Pen-Strep, and sterilized with a 0.22-μm filter). After minced into 1 mm^3^ pieces, tissue was shaken at 37 ^o^C at 150 rpm for 45 min and then centrifuged at 1500 rpm for 3 min. The pellet was suspended in 5 ml ACK buffer and incubated on ice for 5 min to remove red blood cells. 10 ml of DMEM/F12 with 10% FBS was added to neutralize the ACK buffer. Cells were then filtered thought a 40-μm strainer, followed by centrifugation at 1500 rpm for 3 min. The pellet was resuspended in FBS with 10% DMSO, aliquoted into 1 ml per tube, frozen in a Mr. Frosty Freezing Container (Thermo Fisher Scientific) at -80 ^o^C. and then shipped to BGI on dry ice. Upon receiving, the SVF cells were rapidly resuscitated in 37 ^o^C water bath. After sorting with Dead cell removal kit (Miltenyi Biotec), live SVF cells were used for library construction using the Chromium Controller and Chromium Single Cell 3’ Reagent Kit (10X Genomics). The library was sequenced using a BGISEQ-500 instrument.

FASTQ reads were processed and converted to digital gene expression matrices after mapping to the reference genome using the Cell Ranger Single Cell Software Suite (v3.1.0). Seurat (v3.1.5) was used to trim dataset (min.cells = 9, nFeature_RNA > 200, nFeature_RNA < 6380, percent.mt < 5), normalize data (LogNormalize). Highly variable genes were used for principal component analysis, followed by clustering in PCA space using a graph-based clustering approach (dims = 1:10, resolution = 0.5). UMAP and tSNE were then used for two-dimensional visualization of the resulting cluster. Marker genes were identified using the FindAllMarkers function (only.pos = TRUE, min.pct = 0.25, logfc.threshold = 0.25).

**Statistical analyses**

Results are shown as mean ± SEM. The comparisons were carried out using two-tailed unpaired Student’s t-test (Excel) and one-way or two-way ANOVA with indicated post hoc tests with Prism 7 (Graphpad).
