## Supplemental Figures for "Loss of FSTL1-expressing adipocyte progenitors drives the age-related involution of brown adipose tissue"

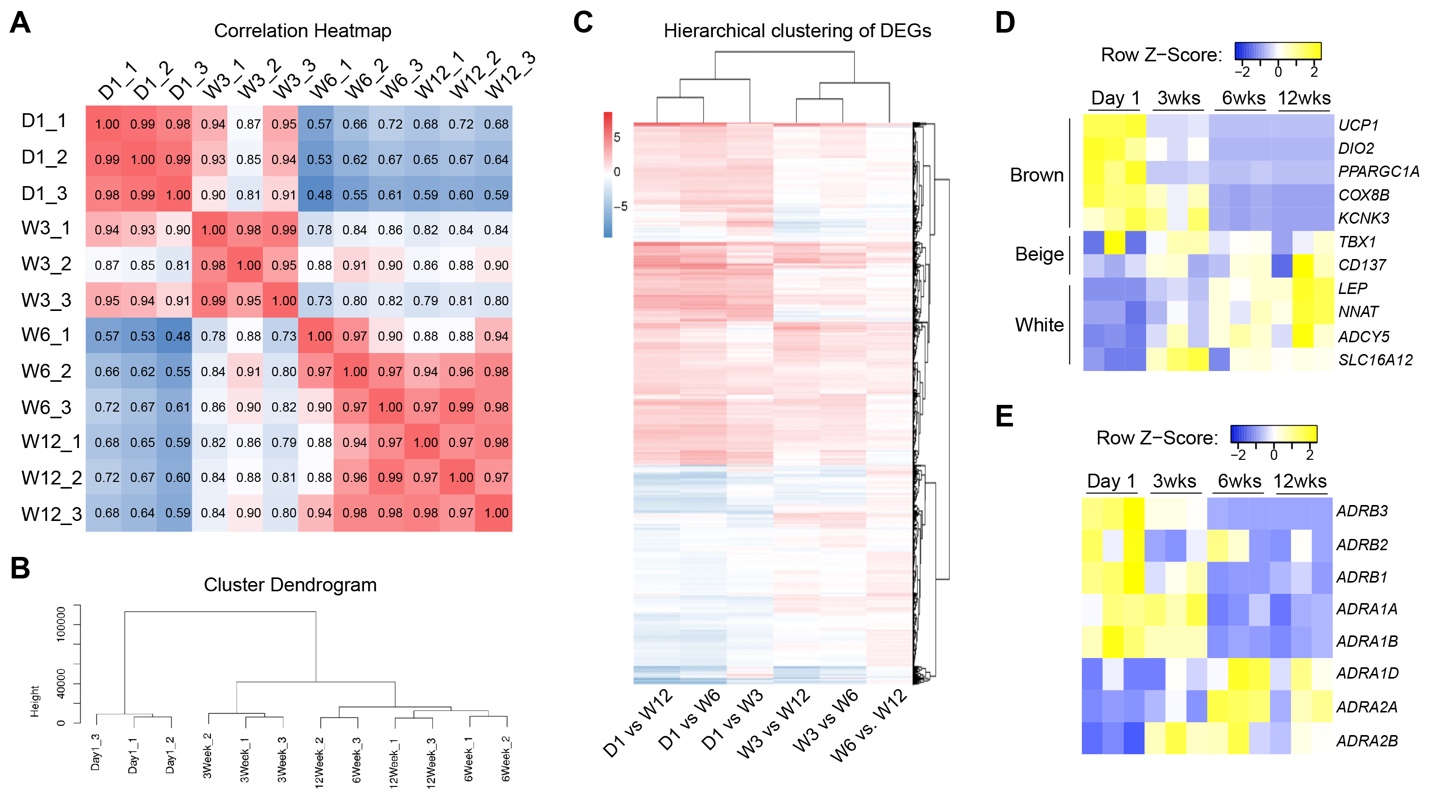

**Figure S1, Related to Figure 1.**

(A, B) Pearson correlation coefficients for all gene expression levels between each two iBAT samples were calculated. A, reflected coefficients in the form of heatmaps. B, hierarchical clustering of all samples by the expression level of all genes

(C) Hierarchical clustering of DEGs. Coloring indicates the log2 of transformed fold change (high: red, low: blue).

(D, E) Expression heatmap of adipocyte marker genes (D) and adrenergic receptor genes (E). Coloring indicates FPKM (fragments per kilobase of exon model per million reads mapped) values (high: yellow, low: blue).

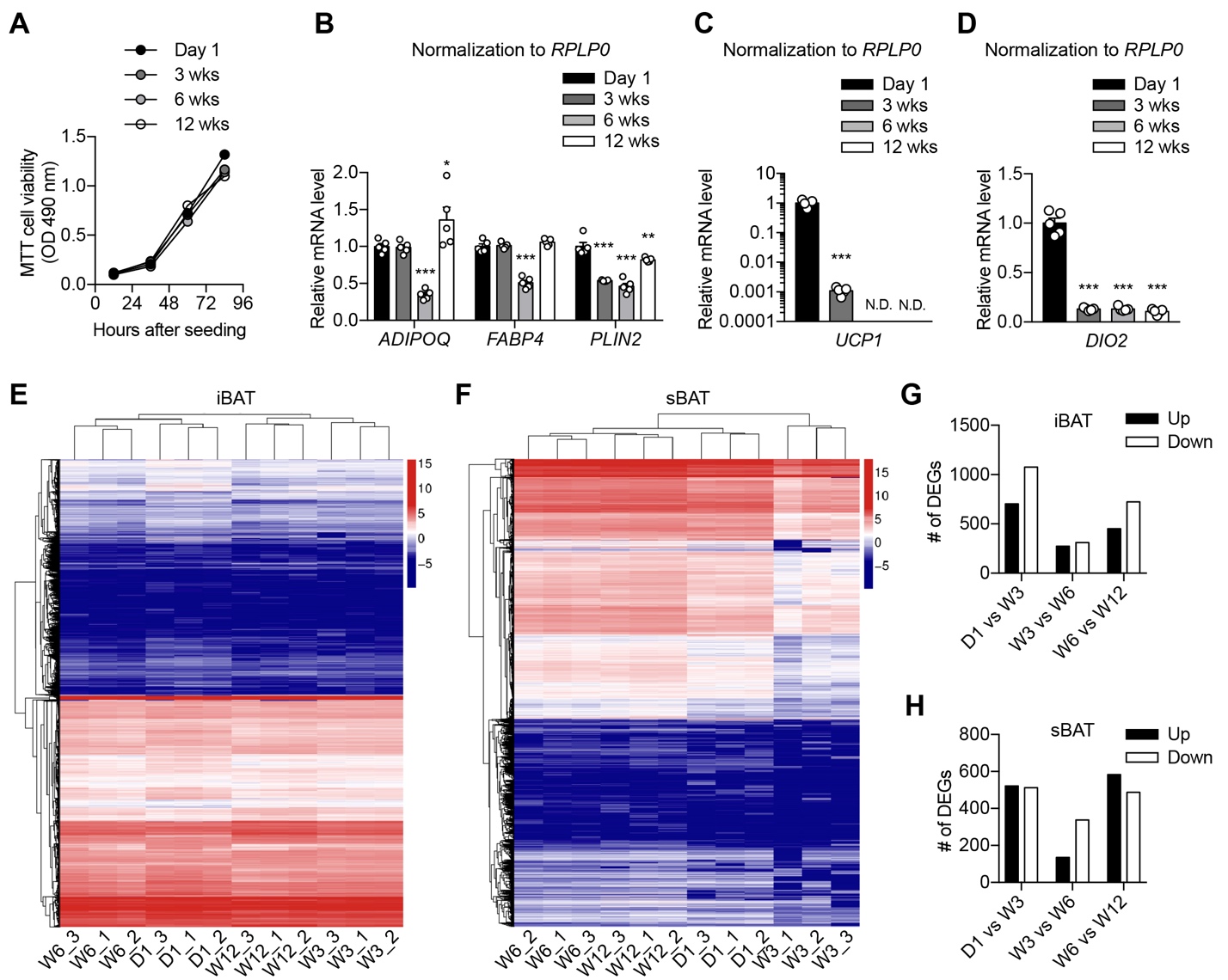

**Figure S2, Related to Figure 2.**

(A) MTT assay showing the proliferation rates of iBAT SVF cells from rabbits at different ages (n = 6).

(B-D) Relative gene expression in differentiated iBAT SVF cells, when normalized to the housekeeping gene *RPLP0* (n = 5).

(E, F) Hierarchical clustering of DEGs in iBAT SVFs (E) and sBAT SVFs (F). Coloring indicates the log2 of transformed fold change (high: red, low: blue).

(G, H) Numbers of differently expressed genes (DEGs) between indicated age groups in iBAT (G) and sBAT (H) SVFs.

Data are presented as mean ± SEM (B-D). *, P < 0.05; **, P < 0.01; ***, P < 0.001 by one-way ANOVA followed with Tukey’s multiple comparison.

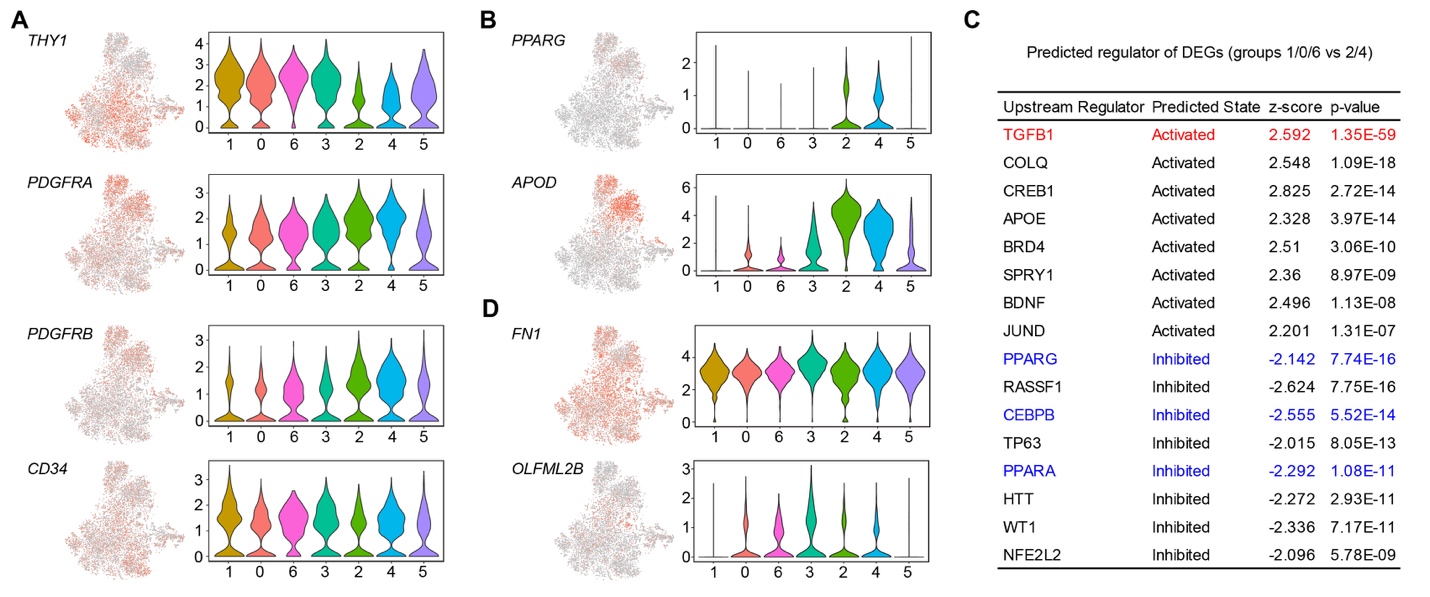

**Figure S3, Related to Figure 3.**

(A, B) tSNE and violin plots showing the expression levels and distribution of mesenchymal (A) and preadipocyte (B) marker genes.

(C) List of IPA-predicted upstream regulators of differentially expressed genes (DEGs) between adipocyte progenitors and committed preadipocytes.

(D) tSNE and violin plots showing the expression levels and distribution of Group 3 marker genes.

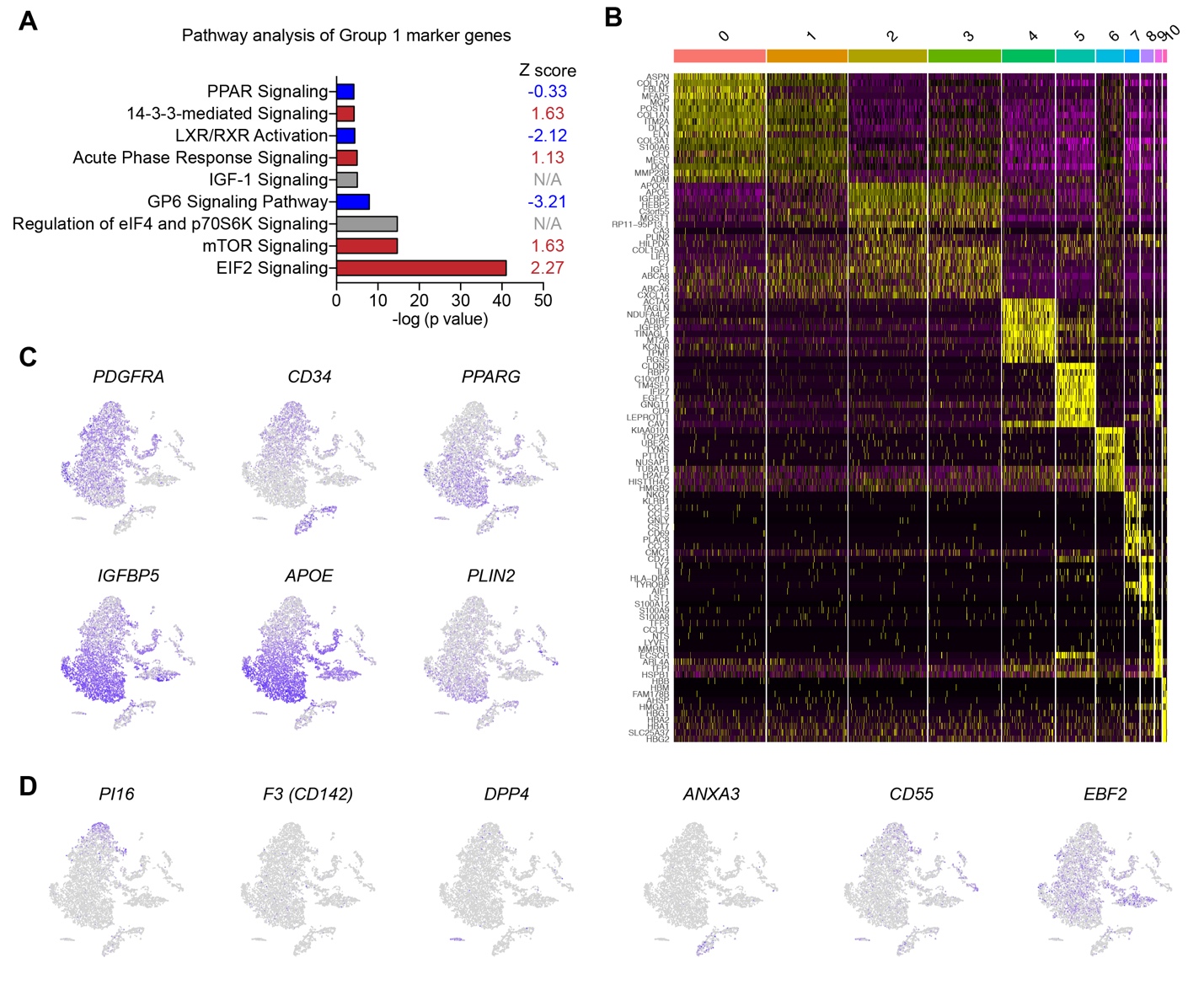

**Figure S4, Related to Figure 4.**

(A) IPA pathway analysis of specific marker genes of Group 1 cells from rabbit iBAT SVF. The Z-score infers the activation state of a putative regulator (red: activation; blue: inhibition).

(B) The expression heatmap of top 25 markers for each cluster identified from human fetal iBAT SVF cells.

(C) tSNE plots of human fetal iBAT SVF cells showing the distribution of marker genes for brown adipocyte progenitors and committed brown preadipocytes.

(D) tSNE plots showing the distribution of white adipocyte progenitor markers in human fetal iBAT SVF cells.

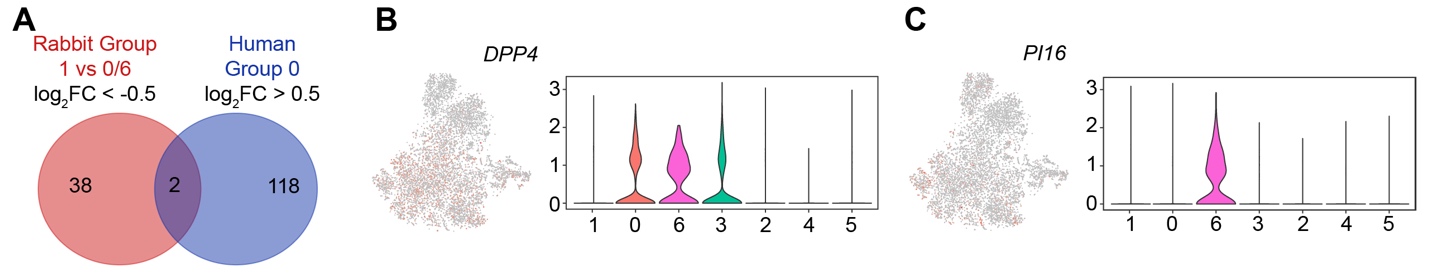

**Figure S5, Related to Figure 5.**

(A) Venn diagram showing little overlap between group 0/6 adipocyte progenitors of involuted rabbit iBAT and group 0 brown adipocyte progenitors of human fetal iBAT.

(B, C) tSNE and violin plots showing the expression levels and distribution of white adipocyte progenitor marker *DPP4* (C) and *PI16* (D) in rabbit iBAT SVF cells.

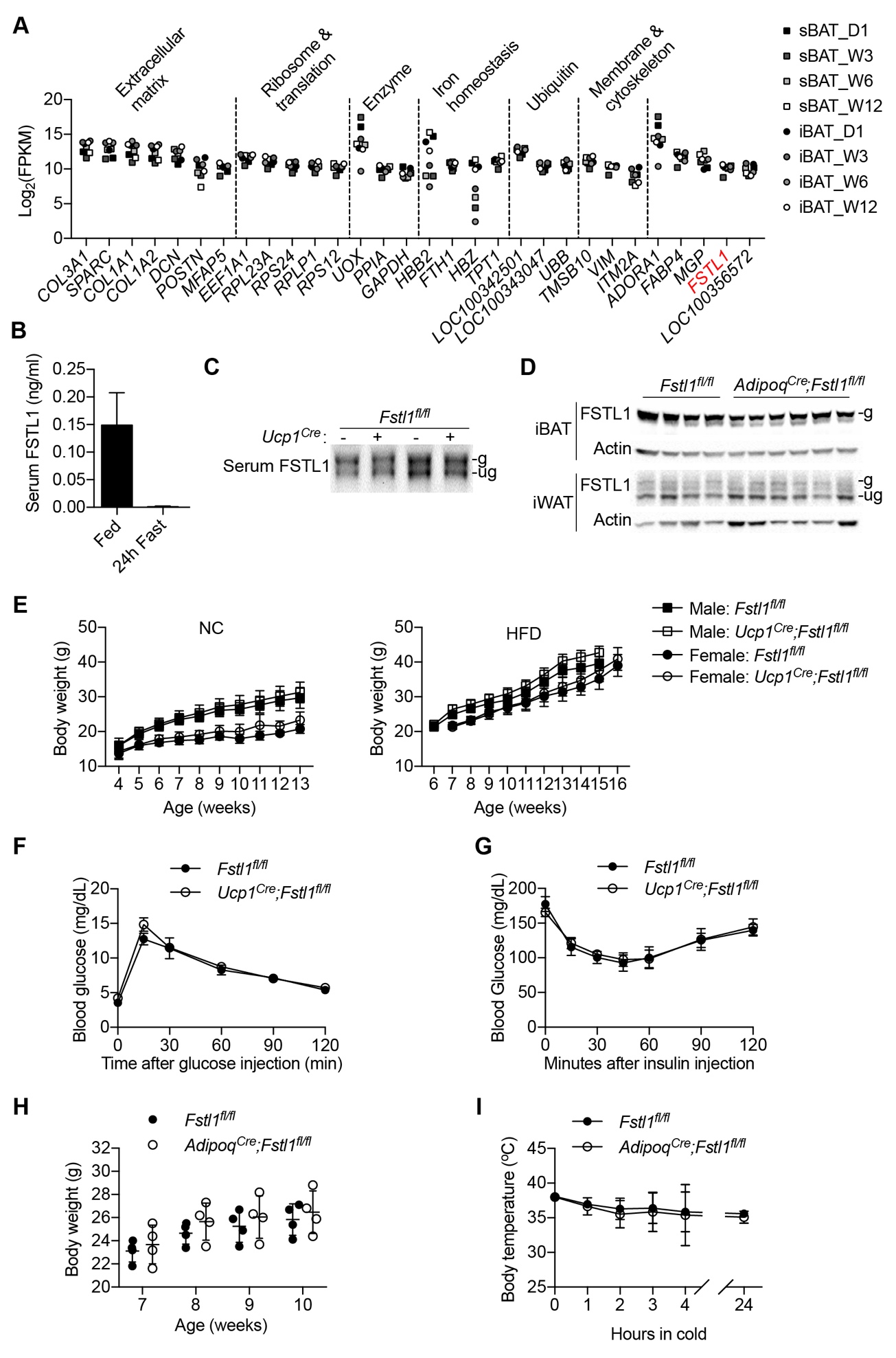

**Figure S6, Related to Figure 6.**

(A) Top 30 expressed genes in total sBAT and iBAT depots of rabbits, based on FPKM values from RNA-seq.

(B) FSTL1 levels in the serum of mice either fed *ad libitum* or fasted for 24 h determined by ELISA (n = 5).

(C) FSTL1 levels in the serum of *Fstl1^fl/fl^* and *Ucp1^Cre^;Fstl1^fl/fl^* mice.

(D) FSTL1 protein expression in iBAT and iWAT depots of *Fstl1^fl/fl^* and *Adipoq^Cre^;Fstl1^fl/fl^* mice. “g” and “ug” indicate glycosylated and unglycosylated FSTL1, respectively.

(E) Growth curve of male and female *Fstl1^fl/fl^* and *Ucp1^Cre^;Fstl1^fl/fl^* mice fed with normal chow (NC) or high fat diet (HFD) (n = 5-9).

(F, G) Glucose tolerance test (F) and insulin tolerance test (G) of female *Fstl1^fl/fl^* and *Ucp1^Cre^;Fstl1^fl/fl^* mice (n = 4-6).

(H) Body weight of *Fstl1^fl/fl^* and *Adipoq^Cre^;Fstl1^fl/fl^* male mice fed with normal chow at indicated ages (n = 4).

(I) Core body temperature of *Fstl1^fl/fl^* and *Adipoq^Cre^;Fstl1^fl/fl^* male mice during cold challenge (n = 6).

Data are presented as mean ± SEM.

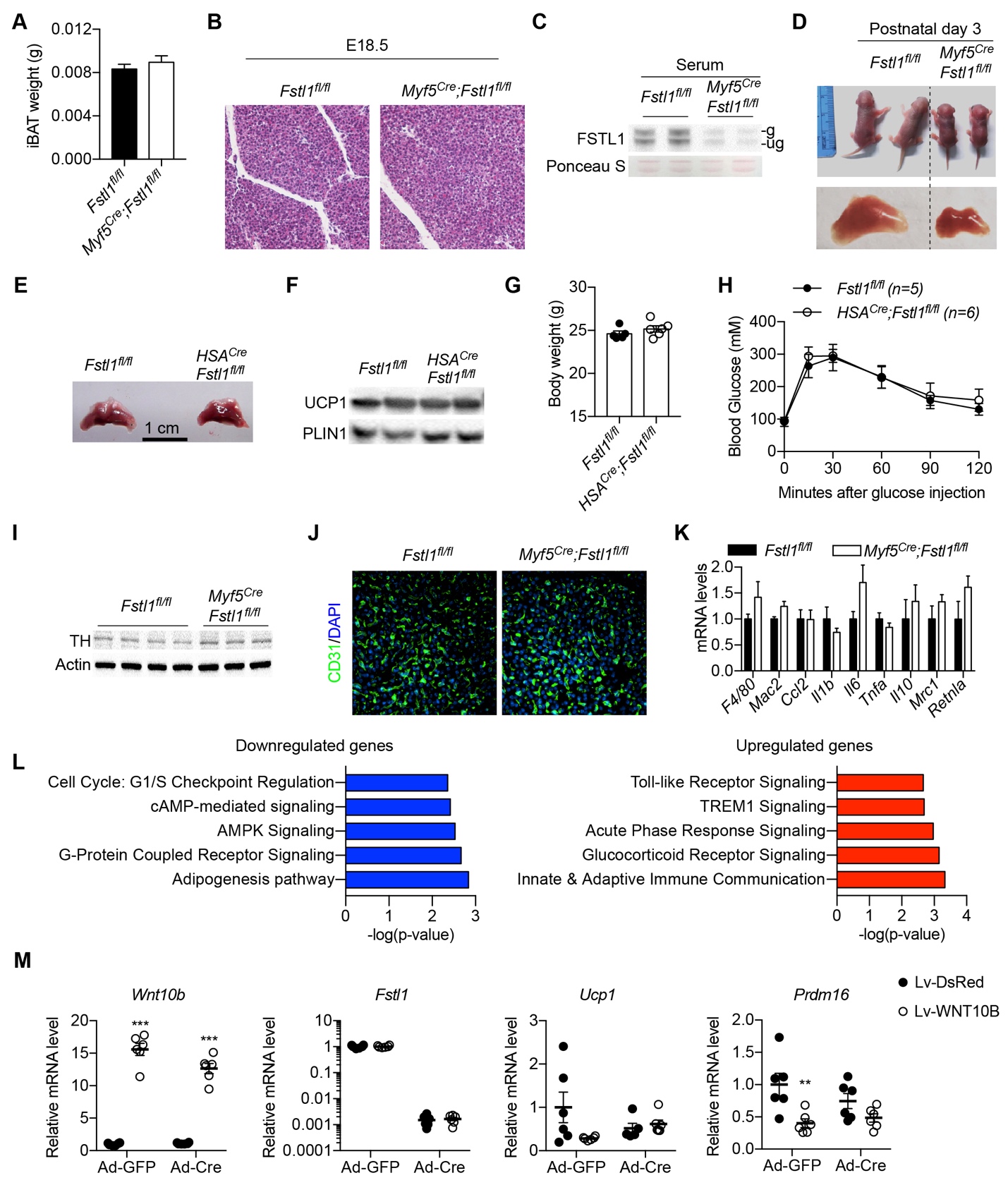

**Figure S7, Related to Figure 7.**

(A, B) Weight (A) and histology (B) of iBAT from control and *Myf5^Cre^;Fstl1^fl/fl^* knockout mice at embryonic day 18.5.

(C) Levels of serum FSTL1 in neonatal control and *Myf5^Cre^;Fstl1^fl/fl^* knockout mice, determined by Western blotting.

(D) Pictures of 3-day-old control and *Myf5^Cre^;Fstl1^fl/fl^* knockout mice housed at RT and the size of iBAT.

(E-H) iBAT morphology (E), UCP1 and PLIN2 protein expression (F), body weight (G), and glucose tolerance (H) of control and *HSA^Cre^;Fstl1^fl/fl^* knockout mice.

(I) Levels of TH protein expression in iBAT of control and *Myf5^Cre^;Fstl1^fl/fl^* knockout mice.

(J) Vascular morphology in iBAT of control and *Myf5^Cre^;Fstl1^fl/fl^* knockout mice, demonstrated by CD31 immunofluorescent staining.

(K) Expression of macrophage marker genes in iBAT of control and *Myf5^Cre^;Fstl1^fl/fl^* knockout mice.

(L) IPA analysis of pathways enriched in downregulated (left) and upregulated (right) genes in *Myf5^Cre^;Fstl1^fl/fl^* knockout iBAT.

(M) Western blotting showing FSTL1 knockdown in *Fstl1^fl/fl^* SVF infected with adenovirus expressing the Cre recombinase.

Data are presented as mean ± SEM. **, p < 0.01; ***, p < 0.001 by two-way ANOVA (M).
